# Modeling and Design of Multi-layered Cylindrical Microcapsules for Intravitreal Controlled Release

**DOI:** 10.64898/2025.12.29.696951

**Authors:** Eduardo A. Chacin Ruiz, Katelyn E. Swindle-Reilly, Ashlee N. Ford Versypt

**Affiliations:** Department of Chemical and Biological Engineering, University at Buffalo, The State University of New York, Buffalo, 14260, NY, USA; Department of Biomedical Engineering, The Ohio State University, Columbus, 43210, OH, USA; William G. Lowrie Department of Chemical and Biomolecular Engineering, The Ohio State University, Columbus, 43210, OH, USA; Department of Ophthalmology and Visual Sciences, The Ohio State University, Columbus, 43210, OH, USA; Department of Biomedical Engineering, University at Buffalo, The State University of New York, Buffalo, 14260, NY, USA; Department of Pharmaceutical Sciences, University at Buffalo, The State University of New York, Buffalo, 14214, NY, USA; Institute for Artificial Intelligence and Data Science, University at Buffalo, The State University of New York, Buffalo, 14260, NY, USA

**Keywords:** polymeric drug delivery, mechanistic drug release model, polycaprolactone, chitosan

## Abstract

Chronic diseases often require repeated oral or local administration, which can compromise patient compliance. In wet age-related macular degeneration (AMD), current therapies rely on intravitreal injections of anti-vascular endothelial growth factor agents every four to six weeks to maintain therapeutic drug levels. Controlled-release drug delivery systems offer a promising alternative by reducing injection frequency and extending drug release. In this study, we developed a continuum diffusion model to describe drug transport through porous polymeric microcapsules, implemented using the finite element method in COMSOL Multiphysics. The case study focused on cylindrical microcapsules fabricated with either a single polycaprolactone (PCL) layer or a bi-layered chitosan–PCL structure, tested at two capsule sizes and three salt leaching concentrations. Bovine serum albumin and bevacizumab were used as model drugs. Parameter estimation was performed using published release data, with a progressive fitting strategy that carried forward parameters from simpler systems into more complex designs. The model reproduced experimental release profiles across formulations and identified key transport parameters governing release dynamics, including porosity, tortuosity, and mass transfer rates. Design exploration revealed that polymer thickness was the dominant factor controlling release, while addition of the chitosan layer moderated the initial burst and extended therapeutic delivery. This framework demonstrates how computational modeling can reduce experimental burden, guide design optimization, and support the development of long-acting intravitreal drug delivery systems to treat wet AMD by linking drug release kinetics to design variables.

## 1 Introduction

Chronic diseases are commonly treated through repeated oral dosing or frequent injections aimed at slowing disease progression, yet such regimens are associated with poor patient compliance [1]. One example is wet age-related macular degeneration (AMD), which requires intravitreal injections every four to six weeks [2] to overcome ocular delivery barriers and maintain therapeutic drug concentrations [3].

Wet AMD is a progressive retinal disorder that primarily affects the macula, a specialized structure in the retina responsible for sharp and central vision [4]. Wet AMD is caused by elevated secretion of vascular endothelial growth factor (VEGF) [5], which promotes pathological angiogenesis. As a result, new and abnormally leaky blood vessels form within the retina [6], allowing fluid to accumulate in the macula. This leakage damages macular photoreceptors [4], ultimately leading to irreversible vision loss.

Because VEGF plays a central role in disease progression, the current standard of care relies on frequent intravitreal injections of anti-VEGF agents [7]. However, to overcome the burden of repeated injections, numerous studies have investigated anti-VEGF-loaded drug delivery systems (DDSs) designed to reduce injection frequency, prolong drug release, and improve adherence to therapy [8–10].

Among the emerging DDS strategies, biodegradable polymer-based microcapsules have gained traction for their ability to encapsulate substantial drug payloads within layered polymeric shells. Building on this concept, Jiang et al. [11] developed cylindrical microcapsules in which drug is loaded into a hollow core, and the caps are sealed by crimping and melting. Two variations of this design were explored: one consisting of a single polycaprolactone (PCL) annulus surrounding the core, and another bilayered annulus composed of chitosan coated by a concentric PCL outer layer. Both chitosan and PCL are biodegradable and biocompatible polymers [12, 13], yet they contribute differently in the release mechanism. Chitosan can enhance retention of negatively charged drugs like bovine serum albumin (BSA) or bevacizumab due to electrostatic interactions [14, 15]. PCL is hydrophobic and degrades slowly, making it suitable for sustained-release applications [16–18]. The combination of these materials offers a promising platform for controlled drug delivery, balancing stability with tunable release kinetics.

Two capsule sizes (inner diameter of 0.260 mm and 1.645 mm) were fabricated in the study by Jiang et al. [11]. The fabrication process involved electrospinning of the polymer fibers onto rotating stainless steel rods of the desired diameter. PCL nanofibers were electrospun from PCL/HEPES solutions at varying ratios (5%, 7.5%, or 10% of HEPES sodium salts), and chitosan was electrospun without salts. Then, the resulting PCL-only or chitosan-PCL microcapsules were sintered to remove surface porosity. Following sintering, capsules were neutralized, washed to remove salts to induce controlled porosity, and vacuum-dried before drug loading and final sealing. The salt concentrations determined three different formulations with their pore diameter distributions (Figure A1), assuming normal distributions around the reported means and standard deviations [11].

Predicting sustained release from polymeric DDSs depends on identifying the parameters that govern drug release kinetics. Yet, these parameters are often not precisely known in advance, and in many cases the dominant release mechanism itself might be uncertain. This uncertainty requires extensive experimental testing to characterize drug release behavior, a process that is both time-consuming and costly. The need for repeated experiments becomes even more important in systems where structural modifications (like varying salt concentrations to tune porosity) produce different formulations with different release dynamics. To reduce this experimental burden, mathematical modeling has emerged as a powerful tool for predicting drug release across a wide parameter space.

Diffusion, swelling, and erosion are typical mechanisms that can control drug release kinetics from biodegradable polymeric DDSs [9, 19–21]. In this work, we focus on diffusion-controlled release, where drug transport is dominated by diffusion through the fluid-filled pores in the polymer(s). Diffusion refers to the transport of individual molecules of a substance (drug) from areas of higher concentration to areas of lower concentration [22]. This process is driven by Brownian motion, leading to homogeneous distribution of the drug in the fluid bulk [23].

There are three types of mechanisms for diffusive mass transport in fluid-filled pores: continuum diffusion, Knudsen diffusion, and surface diffusion [23]. Continuum diffusion governs particle movement driven by Brownian motion and interparticle collisions within the bulk fluid and where boundary effects are negligible. Knudsen diffusion becomes significant when pore diameters approach or fall below the mean free path of particle motion, and collisions with pore walls dominate over intermolecular interactions. As a result, the movement of the different molecules is independent of each other [24]. Surface diffusion arises when particles adsorb onto pore surfaces rather than reflecting off. The adsorbed species migrate only along the solid interface. Unlike bulk diffusion, this movement is confined to the interfacial plane and results in significantly reduced mass transport. In the context of porous DDS formulations, continuum diffusion typically dominates. The relatively large pore dimensions ensure minimal wall interactions, as the particles’ mean free paths remain short in comparison [23]. Therefore, particle trajectories mainly change due to interparticle collisions.

In this study, we addressed the challenge of optimizing intravitreal DDS design by employing computational models, with the dual aim of reducing the cost and time associated with experimental design and providing guidance for future experimental efforts. We developed a continuum diffusion model to describe drug release from fluid-filled pores in cylindrical DDSs, implemented using the finite element method in COM-SOL. As a case study, we applied the model to PCL and chitosan-PCL microcapsules of two different sizes for the release bovine serum albumin (BSA) and bevacizumab [11]. Parameter estimation was performed using published experimental data, with the strategy of carrying forward fitted parameters from earlier steps into subsequent analyses. This approach allowed us to evaluate whether minimal parameter adjustments were sufficient to reproduce experimental release profiles and to test the extent to which data from one experimental set could inform another. Finally, the estimated parameters were used to explore the influence of the design variables: thickness of polymeric layers and ratio of the polymers. This framework supports model-informed development of intravitreal DDSs to meet specific therapeutic thresholds.

## 2 Methods

### 2.1 Geometry

The DDS considered here is a microcapsule consisting of a cylindrical core into which the drug is initially loaded and an outer coating composed of one or two polymeric layers (Figure 1). The cylinder is characterized by its height *H*_*cylinder*_, the core radius *R*_*core*_, and the radii of the first and second polymeric layers 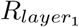 and 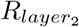, respectively. To simplify the analysis, radial symmetry about the capsule’s central axis is assumed. The specific geometric parameters employed in the model are listed in Table 1.

**Table 1.** Model parameters (obtained from Jiang et al. [11]) used for the microcapsule geometry.

| Small microcapsules |  |  |  |
| --- | --- | --- | --- |
| Definition | Parameter | Value | Units |
| Microcapsule height | $H_c$ | 13 | mm |
| Core radius | $R_{core}$ | 0.13 | mm |
| PCL radius (mono-layered) | $R_{layer_1}$ | 0.22 | mm |
| Chitosan radius (bi-layered) | $R_{layer_1}$ | 0.1525 | mm |
| PCL radius (bi-layered) | $R_{layer_2}$ | 0.22 | mm |
| Large microcapsules |  |  |  |
| Definition | Parameter | Value | Units |
| Microcapsule height | $H_c$ | 13 | mm |
| Core radius | $R_{core}$ | 0.8225 | mm |
| PCL radius (mono-layered) | $R_{layer_1}$ | 0.9125 | mm |
| Chitosan radius (bi-layered) | $R_{layer_1}$ | 0.845 | mm |
| PCL radius (bi-layered) | $R_{layer_2}$ | 0.9125 | mm |

**Fig. 1.**
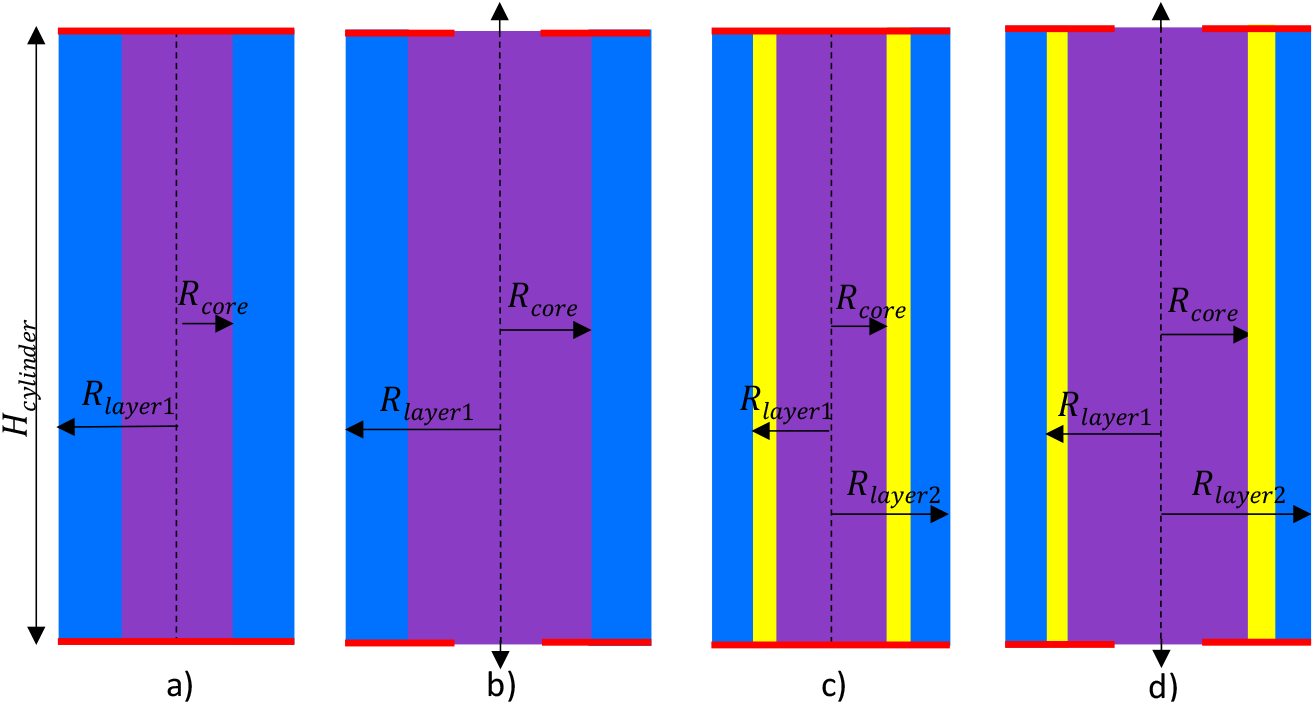
Two-dimensional cylindrical cross-section of drug-loaded microcapsules. a) A mono-layered microcapsule with a small core. b) A mono-layered microcapsule with a large core. c) A bi-layered microcapsule with a small core. d) A bi-layered microcapsule with a large core. A drug is loaded only in the cylindrical core. Purple: homogeneously distributed drug within the core. Blue and yellow: different polymeric layers. *H*: height. *R*: radius. Closed caps (red lines) represent perfect sealing. Open caps (gaps between red lines) allow partial drug release. Note: the microcapsules are not illustrated to scale.

### 2.2 Mathematical model

Drug release from slowly eroding DDSs can be modeled as a diffusion-controlled release mechanism when the time scale for diffusion is faster than the erosion. We assumed that drug diffusion through the porous DDS occurs predominantly through the fluid-filled pores. The mass balance equation for transport in a saturated porous medium assuming no accumulation in the solid phase, no convection (and therefore no dispersion), and no chemical reaction is

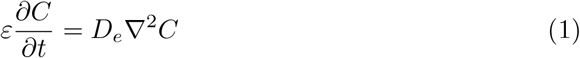

where *ε* is the effective DDS porosity, *C* refers to drug concentration, ∇^2^ is the Laplacian operator, *t* is time, and *D*_*e*_ is an effective diffusion coefficient.

Effective diffusion coefficients can be described at a macroscale DDS level as a function of the DDS porosity and the geometry of the pores as

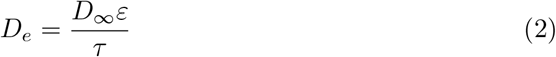

where *D*_∞_ is the drug diffusion coefficient at infinite dilution, and *τ* is the tortuosity. Two important factors cause a reduction in the diffusion coefficient of the drug in a porous DDS: *ε* and *τ* . Since diffusion happens in the fluid-filled pores, the cross-sectional area available for diffusion is lowered by a factor of the porosity *ε*, which is defined as

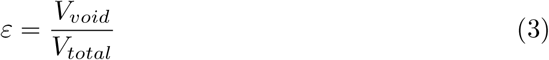

where *V*_*void*_ is the void volume contained in the porous medium and occupied by the fluid, and *V*_*total*_ is the total volume of the same medium. Moreover, since the pores are not straight, the drug needs to diffuse through a longer distance than the actual thickness of the DDS. Therefore, the effective diffusion coefficient is reduced by a factor of the DDS tortuosity *τ*, which is defined as

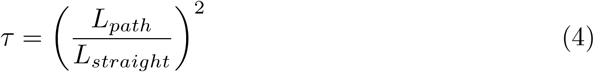

where *L*_*path*_ is the shortest path (measured through connected pores) between any two points of the fluid in a porous medium, and *L*_*straight*_ is the straight-line distance between the same two points. However, directly measuring *L*_*path*_ for determining tortuosity is a non-trivial problem; therefore, simplified correlations have been proposed in the literature [25] to relate tortuosity and porosity. One such is the Bruggeman model [26], which assumes that obstructions to transport are either spheres or cylinders and is based on simple diffusion-controlled transport in an isotropic medium [25]. For cylinders, the approximation is

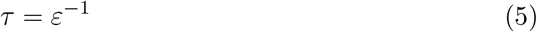

Substituting Equation (5) into Equation (2) yields

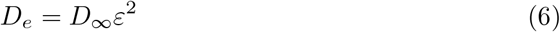

which describes the effective diffusivity as a function of porosity.

The DDS is modeled as a right circular cylinder with a central core surrounded by one or two concentric polymeric layers, with drug transport assumed to occur radially outward through these layers. We assume there is no variation in the azimuthal axis *θ* yielding 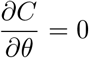. Therefore, Equation (1) becomes

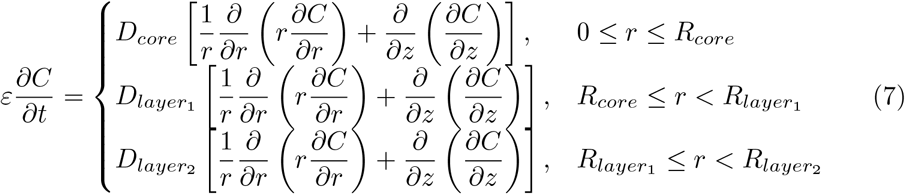

where *D*_*core*_ and 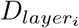 are the effective diffusion coefficients in the core and in each polymeric layer, respectively, *r* is the radial coordinate, and *z* is the height coordinate.

At *r* = 0, a radial symmetry boundary condition is applied:

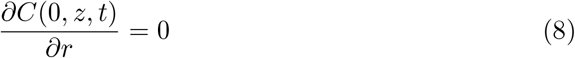

At the interfaces between domains, flux continuity is enforced, and preferential drug partitioning is also permitted. At the first interface,

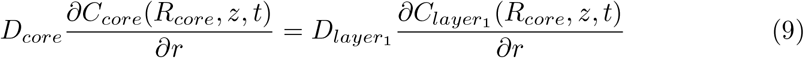

subject to the partition condition

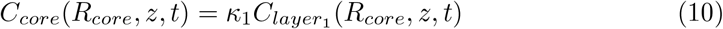

where *C*_*core*_(*R*_*core*_, *z, t*) is the drug concentration in the core at the interface, *κ*_1_ is the partition coefficient between the core and the first layer, and 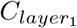 (*R*_*core*_, *z, t*) is the drug concentration in the innermost polymeric layer at the interface.

For a bi-layered microcapsule at the second interface,

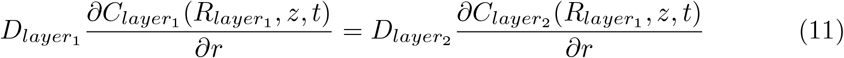

subject to the partition condition

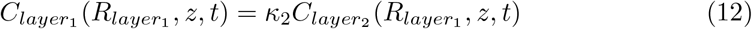

where *κ*_2_ is the partition coefficient between both polymeric layers, and 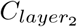 (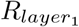, *z, t*) is the drug concentration in the outermost polymeric layer at the interface. Note that we set *κ*_1_ = *κ*_2_ = 1 throughout our analysis because we did not see evidence of partitioning for drugs of interest (BSA and bevacizumab) in our earlier work for bi-layered microspheres [27].

At the external surface of the microcapsule (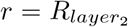 for the bi-layered microcapsule or 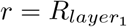 for the mono-layered microcapsule), the perfect sink condition is assumed: *C*(*r, z, t*) = 0.

We assumed perfect sealing of the microcapsules’ caps in the small microcapsules. Therefore, at the top and bottom of the microcapsule (boundaries on the *z* axis marked by red lines in Figure 1a,c), the no-flux boundary condition for drug mass transport is defined by

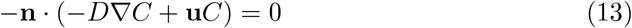

where **n** is the unit normal vector, *D* is the diffusion coefficient of the drug through the medium in contact with the boundary in each layer, and **u** is the fluid velocity inside the microcapsule, which is assumed to be 0.

For the large microcapsules, we assumed that the caps are not perfectly sealed, permitting a finite mass transfer across them. This boundary condition is applied only to the cap area that exceeds the corresponding cap area of the small, perfectly sealed microcapsules. Specifically, a mass transfer resistance boundary condition was imposed on this differential cap area from the centers of the caps (boundaries on the *z* axis between those marked by red lines in Figure 1b,d) and is defined by

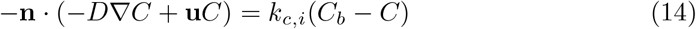

where *k*_*c,i*_ is a mass transfer coefficient for microcapsules composed of PCL-only when *i* = *P* and of chitosan-PCL when *i* = *C*, and *C*_*b*_ is the bulk concentration in the surrounding exterior domain. The mass transfer resistance was estimated separately for the large PCL-only (*k*_*c,P*_ ) and large chitosan–PCL (*k*_*c,C*_) microcapsules, under the assumption of *C*_*b*_ = 0.

The initial drug distribution is defined as

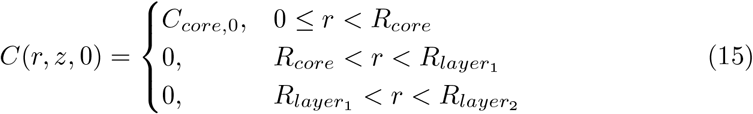

where *C*_*core*,0_ = 1 arbitrary units (a.u.) represents the normalized concentration in the core for all values of *z* immediately after burst release. Burst release is assumed to occur instantaneously at *t* = 0 as it was not observed in the data [11].

Since cumulative release is the experimental metric reported for the DDSs [11], the model prediction is likewise expressed as cumulative release, calculated as

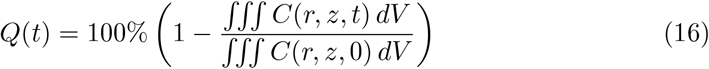

where *Q*(*t*) is the cumulative percentage of drug released as a function of time, *V* is volume, and the last term refers to the fractional amount of drug remaining in the DDS at each time relative to that present initially. The total amount of drug in the microcapsule at any time is calculated as the volume-integral of the spatiotemporal distribution of *C*(*r, z, t*) from the solution of Equations (7) to (15).

The amount of drug released *A*_*rel*_(*t*, Δ*t*) over a given time interval Δ*t* is calculated from the cumulative release profile according to

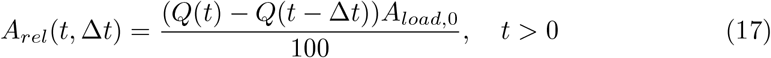

where *A*_*load*,0_ is the initial amount of drug loaded in a microcapsule in moles. Drug-specific molecular weights is used to convert initial amounts from mass to molar units.

The release rate 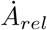 (*t*, Δ*t*) is

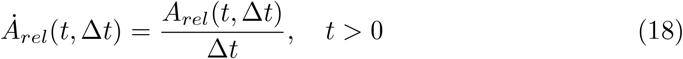

providing the average release rate between two time points separated by Δ*t*. This formulation approximates the derivative of the cumulative release curve, with the highest release rates observed at early times.

### 2.3 Effective diffusivity calculation

Experimental release profiles for all formulations exhibited a bi-phasic behavior, characterized by an initial rapid release over a non-instantaneous timeframe followed by a slower, sustained release (Figure B2). This behavior is consistent with observations reported for other porous DDSs [28–31].

To capture this behavior, the model divides the release process into two distinct phases (or time intervals), each represented by a constant diffusion coefficient. Porosity *ε* was assumed to remain constant throughout, so temporal changes in effective diffusivity were attributed solely to variations in tortuosity *τ* . In this framework, the first phase corresponds to the rapid release of the drug located closer to the capsule surface where tortuosity is lower. The second phase reflects slower release of the drug positioned deeper within the capsule where a longer and more complex diffusion path results in higher tortuosity. For PCL, the effective diffusivity is defined as

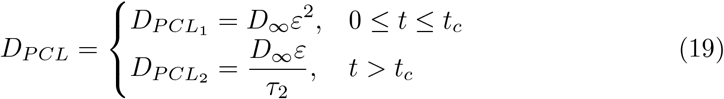

where 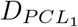 is the drug diffusivity in PCL during the first release phase, 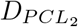 is the drug diffusivity in PCL during the second release phase, *t*_*c*_ is the critical time at which the phase transition happens, and *τ*_2_ is the tortuosity in PCL during the second release phase. For chitosan, the effective diffusivity was scaled relative to that of PCL by

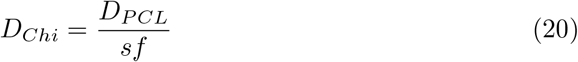

where *sf* is a scaling factor that accounts for reduced transport through the chitosan matrix. Note that 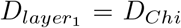 and 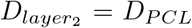 for the bi-layered DDS and 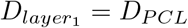 for the mono-layered DDS.

### 2.4 Numerical methods

Equations (7) to (15) and Equations (19) to (20) were solved numerically using the finite element method implemented in COMSOL Multiphysics 6.2. Leveraging axial symmetry, the problem geometry was reduced to a two-dimensional (2D) axisymmetric representation. The computational domain was discretized with a triangular mesh, with element sizes ranging from 2.4 × 10^−5^ to 0.1 mm as in related finite element models [27, 32].

Drug transport in the DDS was modeled using the Transport of Diluted Species interface for the core region, and the Transport of Diluted Species in Porous Media interface for the polymeric coating(s). At the core–polymer and polymer–polymer interfaces, a partition Condition boundary condition was applied. The cumulative release expression (Equation (16)) was solved using the Non-local coupling: integration feature to compute the volume integral of the spatially dependent concentration profile across the microcapsule.

### 2.5 Parameter estimation

The parameter estimation was performed under a progressive fitting approach (Figure 2) starting from the simplest design, the small mono-layered PCL-only micro-capsule. For this microcapsule, the parameters *p* = {porosity *ε* and the final tortuosity *τ*_2_} for Equation (19) were estimated while holding the critical time *t*_*c*_ constant, and 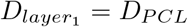. These values of *ε* and *τ*_2_ were then fixed and carried forward into all subsequent parameter estimation steps. For the large mono-layered PCL-only micro-capsules, the parameter set was *p* = {mass transfer rate *k*_*c,P*_ }, which was estimated at a newly defined *t*_*c*_. The choice of *t*_*c*_ was based on experimental measurement points reported in the dataset, ensuring consistency between model assumptions and available observations.

**Fig. 2.**
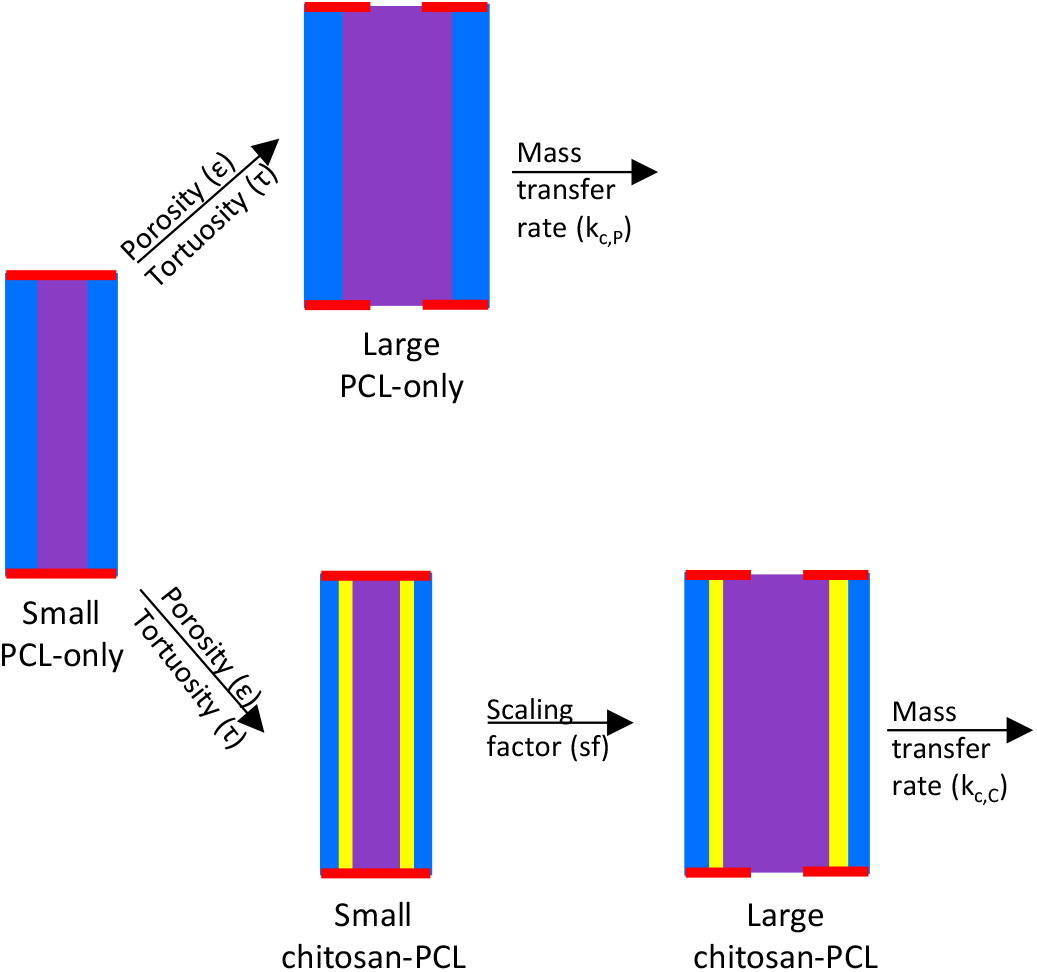
Progressive parameter fitting approach. The arrows denote the parameters carried forward from one case to the next. PCL: polycaprolactone. Note: the microcapsules are not illustrated to scale.

For the small bi-layered microcapsules, the intermediate polymeric layer was expected to slow down drug release. Therefore, the effective diffusivity of 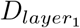 was approximated by dividing the effective diffusivity of the outer layer 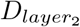 by a new parameter *p* = {scaling factor *sf* }. This parameter was estimated while maintaining the same *t*_*c*_ used for the small mono-layered microcapsules. For the large bi-layered microcapsules, we re-estimated *p* = {mass transfer rate *k*_*c,C*_}. This estimation was performed at the same fixed *t*_*c*_ as for the large mono-layered microcapsules.

Parameter estimation was carried out by minimizing an ordinary least-squares objective function, defined as the sum of squared differences between the model predictions and the experimental observations, for the set of parameters *p*:

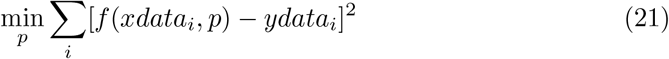

where *f* (*xdata*_*i*_, *p*) represents the model output evaluated at the input values *xdata*_*i*_ corresponding to experimental measurements *ydata*_*i*_, with *i* indexing discrete data points. The data were obtained from Jiang et al. [11], and the model outputs of interest were the cumulative release values defined in Equation (16), evaluated at times *xdata*_*i*_ when the experimental cumulative drug release measurements *ydata*_*i*_ were obtained. Parameter estimation was performed separately for each microcapsule formulation for BSA and bevacizumab, with the minimized sum of squared residuals (Equation (21)) reported as the “error value”.

Parameter estimation for the set of model parameters *p* was performed in COMSOL using the optimization module’s built-in global least-squares objective function. Each parameter was normalized by its respective scaling term so that the midpoint of the parameter range was set to unity. The optimization routine was executed with a convergence tolerance of 0.001 and capped at 1000 function evaluations.

A multi-start strategy was employed to reduce the likelihood of convergence to local minima. Fifty initial parameter sets were generated using Latin hypercube sampling within the bounds specified in Table 2, with uniform sampling applied across the logarithm of each parameter. Parameter sets yielding error values within 5% of the overall minimum value obtained were classified as acceptable. The model output with the average of all acceptable parameter sets did not differ visually from the output obtained with the single parameter set corresponding to the lowest error (Figures C3 to C8 for the PCL-only microcapsules). After comparison, the averaged parameters were selected for all subsequent model simulations.

**Table 2.** Limits used in the multi-start parameter estimation.

| Parameter | Lower limit | Upper limit | Units |
| --- | --- | --- | --- |
| $\varepsilon$ | $1 \times 10^{-3}$ | $1 \times 10^{-1}$ | - |
| $\tau_2$ | $1 \times 10^2$ | $1 \times 10^4$ | - |
| $k_{c,i}$ | $1 \times 10^{-10}$ | $1 \times 10^{-6}$ | cm/s |
| $sf$ | $1 \times 10^0$ | $1 \times 10^2$ | - |

$\varepsilon$ : effective porosity in the polycaprolactone (PCL) layer. $\tau_2$ : tortuosity in the PCL layer during the second release phase. $k_{c,i}$ : mass transfer rate from the caps of the microcapsule, the $i$ denotes microcapsules of PCL-only for $i = P$ and chitosan-PCL for $i = C$ . $sf$ : scaling factor.

Additionally, to provide an absolute measure of the average prediction error in the same units as the dependent variable, the root mean squared error *RMSE* was calculated as

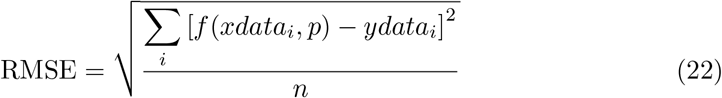

where *n* is the number of observations.

### 2.6 Uncertainty quantification of estimated parameters

Confidence intervals of the estimated parameters were computed from the Levenberg-Marquardt solver from the optimization module in COMSOL [33]. The procedure begins with the calculation of the covariance matrix for the parameter set *p*:

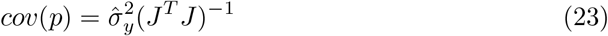

where 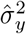 is the variance of the random errors and

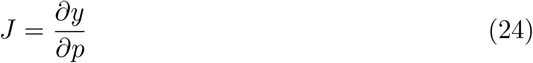

refers to the sensitivity of the simulated measurements *y* to the estimated parameters *p*, which is linearly approximated in COMSOL. The diagonal elements of *cov*(*p*) correspond to the variances of the individual parameters, and their square roots provide the standard deviations *σ*_*j*_. Using these standard deviations, the 95% confidence interval for each parameter *θ*_*j*_ is expressed as

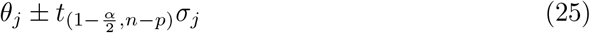

where *n* is the number of data points, *p* is the number of parameters estimated, and *α* is the significance level (*α* = 0.05 for a 95% confidence interval). The term 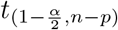 refers to the two-sided *t*-statistic with *n* − *p* degrees of freedom.

### 2.7 Design exploration strategy

Computational models were employed to evaluate the influence of design aspects on bevacizumab release for the PCL-only and chitosan-PCL microcapsules. Release rates were calculated using Equations (17) and (18). The device size here considered was constrained by current ophthalmic standards, specifically the Ozurdex® dexamethasone implant, which is approximately 6 mm long with a diameter of 0.46 mm. For our design exploration simulations, we used this length as the microcapsule height *H*_*c*_ and fixed the microcapsule outer radius at 0.23 mm, which is also very similar to the outer radius for the small microcapsules tested in Jiang et al. [11] and studied here (Table 1).

All design exploration simulations used an initial bevacizumab core concentration *C*_*core*,0_ of 0.75 mg/µL. This loading exceeds that of existing wet AMD injection treatments, such as Vabysmo® (faricimab-svoa, 0.12 mg/µL), but remains below the maximum concentration tested experimentally (1 mg/µL) by Jiang et al. [11]. The 0.75 mg/µL concentration was chosen to compensate for the limited device volume (up to ∼1 µL) compared with the typical 50 µL delivered by intravitreal injections. For each design configuration, the initial drug amount was determined by calculating the core volume and multiplying it by *C*_*core*,0_.

Two design thresholds were established for evaluating the microcapsules. The first was a minimum release rate of 1 µg/day, consistent with prior estimates for ranibizumab, where retinal anti-VEGF production rates of 1.1–2.6 µg/day were considered sufficient to prevent neovascularization [34], and for bevacizumab used in our earlier work [27]. The second threshold was a cumulative drug release of 90%, ensuring that a substantial fraction of the drug is delivered and reflecting practical release scenarios [32].

We assumed that the effective diffusivities in the PCL and chitosan layers remain constant within the design space at the values estimated for the small microcapsules because there are only minor changes in thicknesses. For the PCL-only microcapsules, the outer PCL radius 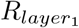 was fixed at 0.23 mm, while the core radius *R*_*core*_ was systematically increased from its baseline value for alternative configurations. For the chitosan-PCL microcapsules, the outer PCL layer 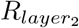 was likewise fixed at 0.23 mm, and two design scenarios were explored: (i) varying the overall polymeric thickness while maintaining a constant PCL-to-chitosan ratio, and (ii) varying the PCL-to-chitosan ratio while holding the total polymeric thickness constant. The thickness of each polymeric annulus is denoted by the symbol *δ* with a subscript corresponding to the polymer (Chi for chitosan or PCL for polycaprolactone).

## 3 Results

### 3.1 Parameter estimation

A preliminary parameter estimation was conducted to identify an appropriate critical time *t*_*c*_ across the three salt leaching concentrations. We performed 50 multi-start parameter estimations at fixed critical times (*t*_*c*_ = 3, 7, and 14 days) for the small PCL-only microcapsules and compared the resulting error values across the 5%, 7.5%, and 10% formulations (Figures C3 to C8 and Table C1). For BSA release, no single critical time consistently outperformed the others across all formulations. We selected *t*_*c*_ = 7 days as the critical time for BSA as it provided acceptable results for all three formulations. In contrast, bevacizumab release consistently showed the lowest errors at *t*_*c*_ = 3 days.

To calculate the parameters following the multi-start approach described in Section 2.5 across all formulations, an arbitrary cut-off threshold was defined as a 5% increase in error relative to the minimum error obtained. Estimated parameters from all simulations with error values below this threshold were considered the acceptable set. The average of each parameter in the acceptable set is summarized alongside its confidence interval in Tables 3 and 4 for BSA and bevacizumab, respectively. The comparison of the parameters with the lowest error values and those in the acceptable set is detailed in Table C1.

**Table 3.** Estimated parameters after fitting the model to data [35] for BSA release from different formulations. The model parameters were obtained by averaging the parameters obtained from all the optimization runs that achieved an error within 5% of the minimum error after 50 multi-start attempts. BSA: bovine serum albumin. *PCL*: polycaprolactone. *t*_*c*_: critical time for phase transition. *ε*: PCL porosity. 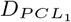 drug diffusivity in PCL during the first release phase. *τ*_2_: PCL tortuosity during the second release phase. 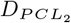 drug diffusivity in PCL during the second release phase. *sf* : scaling factor. 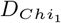 drug diffusivity in chitosan during the first release phase. 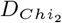 drug diffusivity in chitosan during the second release phase. *k*_*c,P*_ : mass transfer rate in PCL-only microcapsules. *k*_*c,C*_ : mass transfer rate in chitosan-PCL microcapsules. S. L.: salt leaching. C. I.: confidence interval. Error value: sum of squared residuals. RMSE: root mean squared error. *: Calculated values.

| Small PCL-only microcapsules ($t_c = 7$ days) | | | | | | |
| --- | --- | --- | --- | --- | --- | --- |
| S. L. (%) | $\varepsilon \pm 95\%$ C. I. | $DPCL_1$ (cm<sup>2</sup>/s)* | $\tau_2 \pm 95\%$ | $DPCL_2$ (cm<sup>2</sup>/s)* | Error value | RMSE |
| 5 | $(6.73 \pm 0.52) \times 10^{-3}$ | $3.81 \times 10^{-11}$ | $852.7 \pm 193$ | $6.64 \times 10^{-12}$ | 186.5 | 3.31 |
| 7.5 | $(8.14 \pm 0.97) \times 10^{-3}$ | $5.57 \times 10^{-11}$ | $968.5 \pm 490$ | $7.07 \times 10^{-12}$ | 476.6 | 5.83 |
| 10 | $(8.44 \pm 0.45) \times 10^{-3}$ | $5.99 \times 10^{-11}$ | $450.5 \pm 123$ | $1.58 \times 10^{-11}$ | 77.07 | 2.13 |
| Small chitosan-PCL microcapsules ($t_c = 7$ days) | | | | | | |
| S. L. (%) | $sf \pm 95\%$ C. I. | $DChi_1$ (cm<sup>2</sup>/s)* | $DChi_2$ (cm<sup>2</sup>/s)* | Error value | RMSE | |
| 5 | $7.80 \pm 0.42$ | $4.88 \times 10^{-12}$ | $8.51 \times 10^{-13}$ | 70.28 | 1.98 | |
| 7.5 | $7.85 \pm 0.76$ | $7.10 \times 10^{-12}$ | $9.01 \times 10^{-13}$ | 247.2 | 3.71 | |
| 10 | $19.5 \pm 3.77$ | $3.07 \times 10^{-12}$ | $8.10 \times 10^{-13}$ | 1288 | 7.83 | |
| Large PCL-only microcapsules ($t_c = 28$ days) | | | | | | |
| S. L. (%) | $k_{c,P} \pm 95\%$ C. I. (cm/s) | Error value | RMSE | | | |
| 5 | $(2.66 \pm 0.41) \times 10^{-8}$ | 553.3 | 4.53 | | | |
| 7.5 | $(7.82 \pm 2.00) \times 10^{-8}$ | 1390 | 7.18 | | | |
| 10 | $(5.18 \pm 1.31) \times 10^{-7}$ | 580.4 | 5.84 | | | |
| Large chitosan-PCL microcapsules ($t_c = 28$ days) | | | | | | |
| S. L. (%) | $k_{c,C} \pm 95\%$ C. I. (cm/s) | Error value | RMSE | | | |
| 5 | $(1.13 \pm 0.09) \times 10^{-8}$ | 79.23 | 1.82 | | | |
| 7.5 | $(8.83 \pm 0.69) \times 10^{-9}$ | 68.65 | 1.59 | | | |
| 10 | $(1.26 \pm 0.09) \times 10^{-8}$ | 154.3 | 2.39 | | | |

**Table 4.**
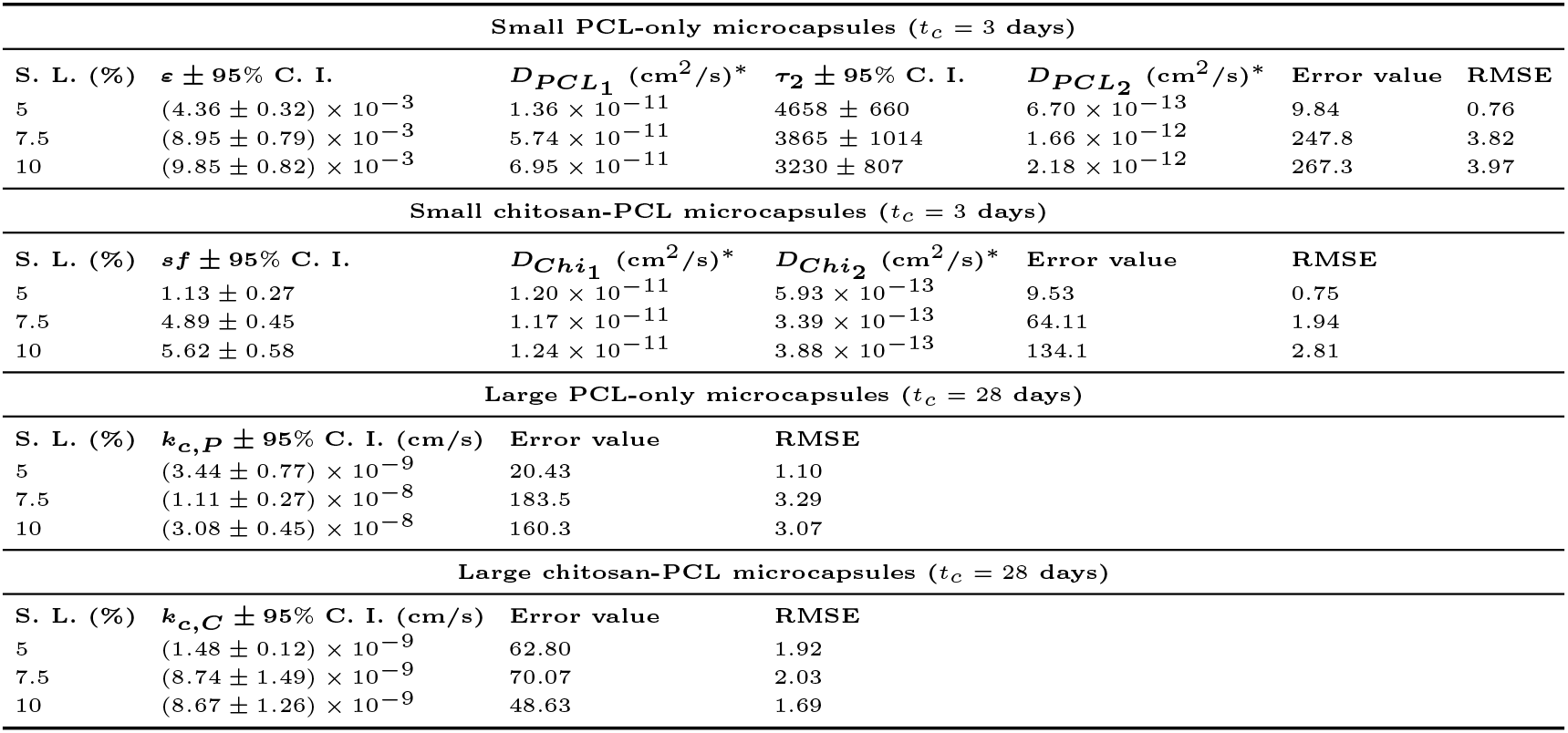
Estimated parameters after fitting the model to data [35] for bevacizumab release from different formulations. The model parameters were obtained by averaging the parameters obtained from all the optimization runs that achieved an error within 5% of the minimum error after 50 multi-start attempts. BSA: bovine serum albumin. *PCL*: polycaprolactone. *t*_*c*_: critical time for phase transition. *ε*: PCL porosity. 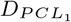 drug diffusivity in PCL during the first release phase. *τ*_2_: PCL tortuosity during the second release phase. 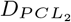 drug diffusivity in PCL during the second release phase. *sf* : scaling factor. 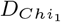 drug diffusivity in chitosan during the first release phase. 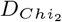 drug diffusivity in chitosan during the second release phase. *k*_*c,P*_ : mass transfer rate in PCL-only microcapsules. *k*_*c,C*_ : mass transfer rate in chitosan-PCL microcapsules. S. L.: salt leaching. C. I.: confidence interval. Error value: sum of squared residuals. RMSE: root mean squared error. *: Calculated values.

| Small PCL-only microcapsules ($t_c = 3$ days) | | | | | | |
| --- | --- | --- | --- | --- | --- | --- |
| S. L. (%) | $\epsilon \pm 95\%$ C. I. | $D_{PCL_1}$ (cm<sup>2</sup>/s)* | $\tau_2 \pm 95\%$ C. I. | $D_{PCL_2}$ (cm<sup>2</sup>/s)* | Error value | RMSE |
| 5 | $(4.36 \pm 0.32) \times 10^{-3}$ | $1.36 \times 10^{-11}$ | $4658 \pm 660$ | $6.70 \times 10^{-13}$ | 9.84 | 0.76 |
| 7.5 | $(8.95 \pm 0.79) \times 10^{-3}$ | $5.74 \times 10^{-11}$ | $3865 \pm 1014$ | $1.66 \times 10^{-12}$ | 247.8 | 3.82 |
| 10 | $(9.85 \pm 0.82) \times 10^{-3}$ | $6.95 \times 10^{-11}$ | $3230 \pm 807$ | $2.18 \times 10^{-12}$ | 267.3 | 3.97 |
| Small chitosan-PCL microcapsules ($t_c = 3$ days) | | | | | | |
| S. L. (%) | $sf \pm 95\%$ C. I. | $D_{Chi_1}$ (cm<sup>2</sup>/s)* | $D_{Chi_2}$ (cm<sup>2</sup>/s)* | Error value | RMSE | |
| 5 | $1.13 \pm 0.27$ | $1.20 \times 10^{-11}$ | $5.93 \times 10^{-13}$ | 9.53 | 0.75 | |
| 7.5 | $4.89 \pm 0.45$ | $1.17 \times 10^{-11}$ | $3.39 \times 10^{-13}$ | 64.11 | 1.94 | |
| 10 | $5.62 \pm 0.58$ | $1.24 \times 10^{-11}$ | $3.88 \times 10^{-13}$ | 134.1 | 2.81 | |
| Large PCL-only microcapsules ($t_c = 28$ days) | | | | | | |
| S. L. (%) | $k_{c,P} \pm 95\%$ C. I. (cm/s) | Error value | RMSE | | | |
| 5 | $(3.44 \pm 0.77) \times 10^{-9}$ | 20.43 | 1.10 | | | |
| 7.5 | $(1.11 \pm 0.27) \times 10^{-8}$ | 183.5 | 3.29 | | | |
| 10 | $(3.08 \pm 0.45) \times 10^{-8}$ | 160.3 | 3.07 | | | |
| Large chitosan-PCL microcapsules ($t_c = 28$ days) | | | | | | |
| S. L. (%) | $k_{c,C} \pm 95\%$ C. I. (cm/s) | Error value | RMSE | | | |
| 5 | $(1.48 \pm 0.12) \times 10^{-9}$ | 62.80 | 1.92 | | | |
| 7.5 | $(8.74 \pm 1.49) \times 10^{-9}$ | 70.07 | 2.03 | | | |
| 10 | $(8.67 \pm 1.26) \times 10^{-9}$ | 48.63 | 1.69 | | | |

The model predictions presented in Figures 3 and 4 were generated using the tabulated parameter values. We also present the same predictions and data focusing on the first 30 days in Figures D9 and D10.

**Fig. 3.**
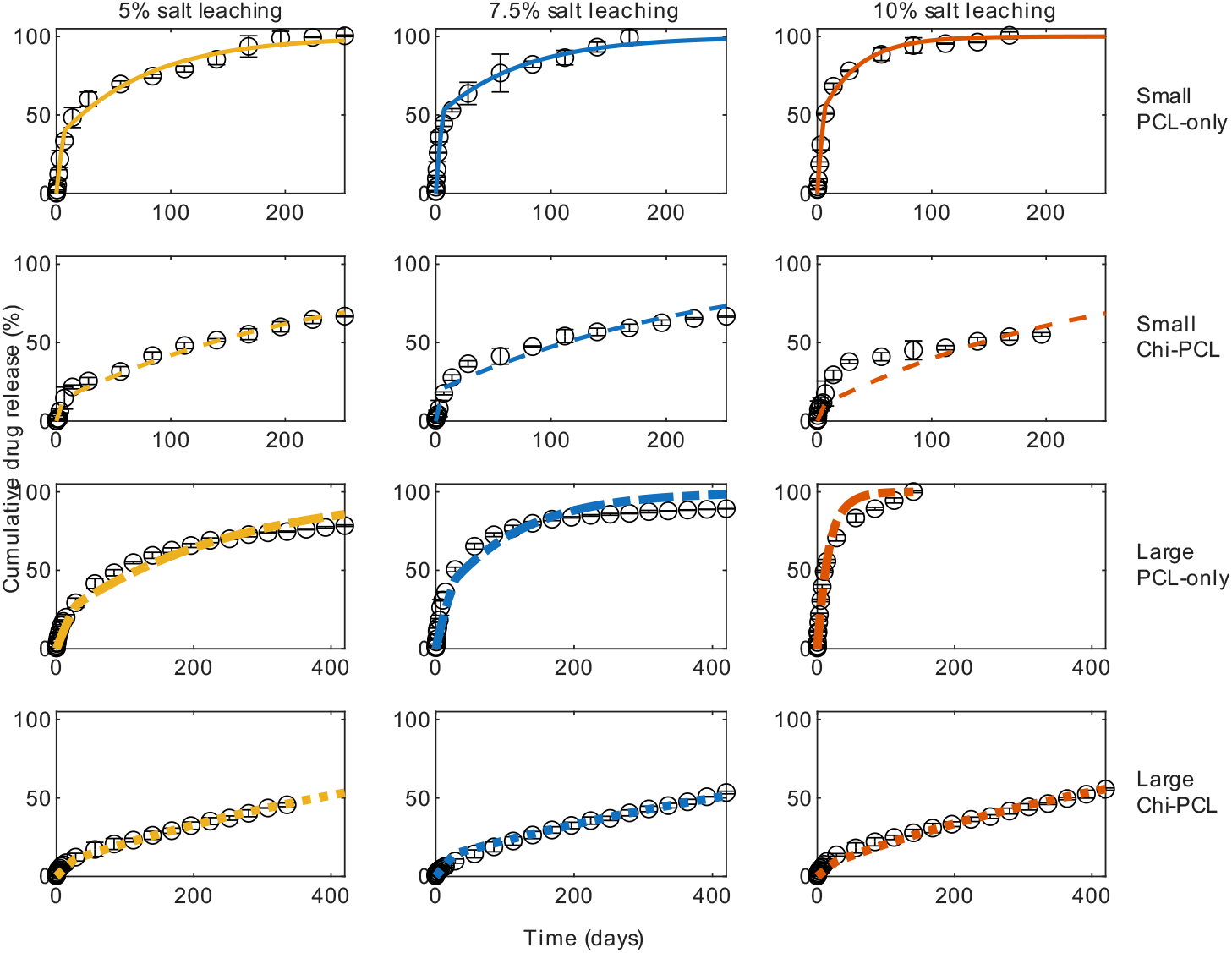
Cumulative BSA release profiles after multi-start parameter estimation with parameters in Table 3 for different microcapsule formulations (salt leaching percentages influencing porosity are labeled across the columns, and size categories for mono- and bi-layered capsules are labeled on the rows). Experimental data from Jiang et al. [11]. BSA: bovine serum albumin. Chi: chitosan. PCL: polycaprolactone.

**Fig. 4.**
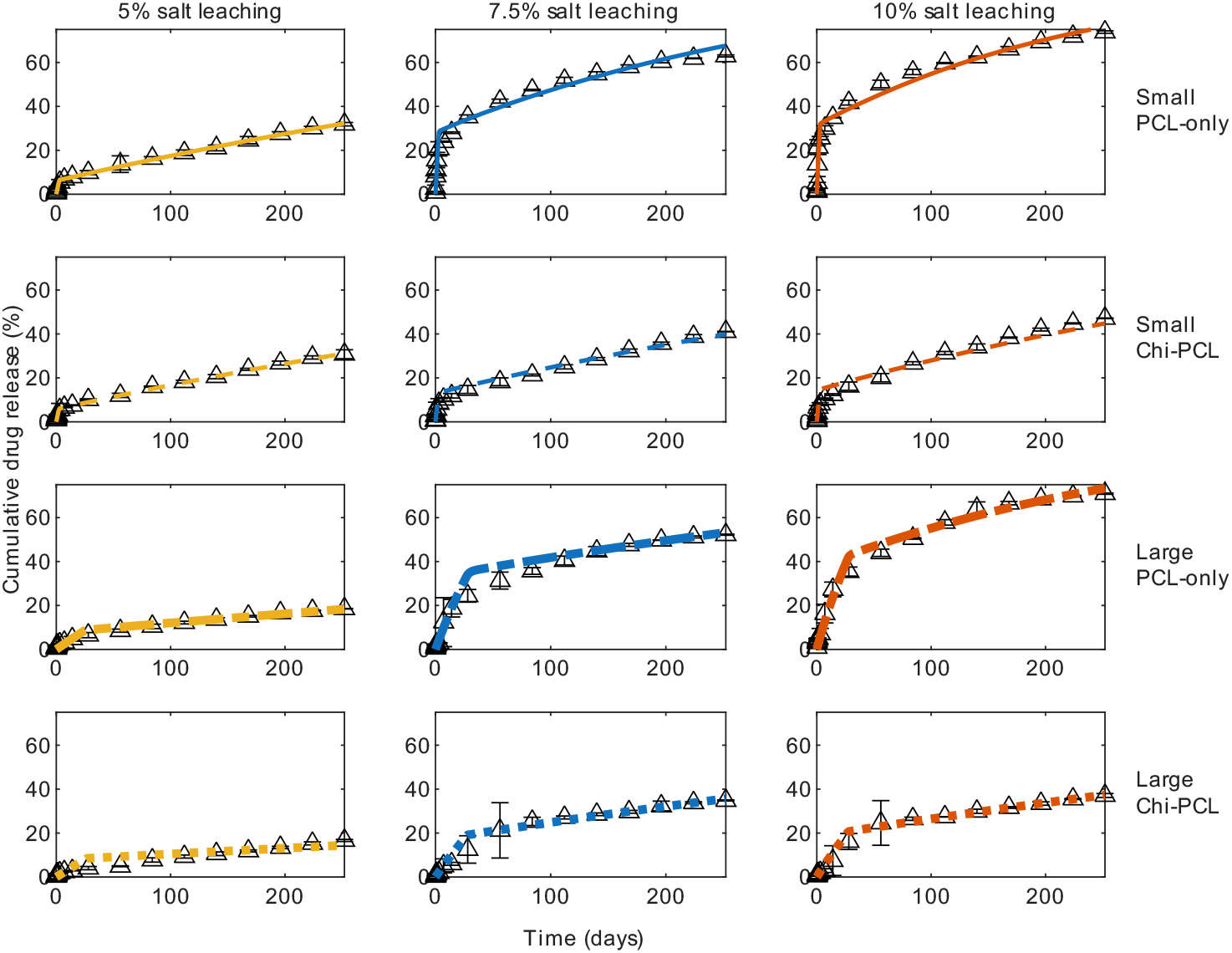
Cumulative bevacizumab release profiles after multi-start parameter estimation with parameters in Table 4 for different microcapsule formulations. Experimental data from Jiang et al. [11]. Chi: chitosan. PCL: polycaprolactone.

For BSA release from small PCL-only microcapsules, the model reproduced the experimental release across all formulations (first row in Figure 3). Some data points’ standard errors exceeded 100% cumulative release, which limited the ability to capture those time points accurately. This could be caused by instrument sensitivity at extended release times or from inaccuracies in calculating the initial drug loading. Increasing the salt concentration led to increased diffusivity (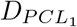 and 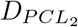), consistent with increased porosities and decreasing tortuosities. An exception was observed in the 7.5% salt leaching formulation, which showed increased tortuosity and the highest RMSE, likely due to the unexpectedly rapid initial release, yielding a high porosity estimate but a longer effective diffusion path during the second release phase.

For the small chitosan–PCL microcapsules, the parameters obtained from the PCL-only microcapsules (*t*_*c*_, *ε*, and *τ*_2_) were used. Because chitosan provides an intermediate barrier layer known to slow the drug release [27], a scaling factor *sf* was introduced to capture this effect. In practice, the chitosan layer was modeled using the PCL parameters divided by *sf* . As shown in the second row in Figure 3, the 5% salt leaching formulation reproduced the release profile accurately, and the 7.5% formulation was reproduced modestly. Both these formulations yielded similar *sf* values, confirming a consistent slowing effect (Table 3). In contrast, the 10% formulation deviated considerably, showing a slower experimental release than expected, and therefore a higher *sf* with the largest RMSE. These differences in *sf* also disrupted the expected trend in chitosan diffusivities 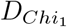 and 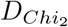, where similar effective diffusivities at each phase were expected. The dependence of cumulative release on salt concentration for salt leaching of chitosan-PCL microcapsules (Figure B2) is not strictly monotonic probably because the 7.5% and 10% formulations yielded very similar pore distributions (Figure A1).

Parameters estimated from the small PCL-only microcapsules were then applied to the large microcapsules for validation. However, the models consistently underpredicted release compared to experimental data (Figure E11, second row). To address this, partial drug release from the end caps of the microcapsules was introduced with a corresponding mass transfer rate *k*_*c,P*_ estimated. The critical time was also extended to *t*_*c*_ = 28 days, reflecting the later phase transition observed in larger microcapsules. As shown in the third row of Figure 3, the 5% salt leaching formulation matched experimental data well, while the 7.5% and 10% formulations deviated to a modest extent. The discrepancy at 7.5% likely arose from the high porosity estimated in the small capsules, whereas the 10% formulation showed unexpectedly faster release in large capsules. Consistent with these trends, Table 3 shows that higher salt concentrations corresponded to higher *k*_*c,P*_ values, reflecting enhanced release with increased salt leaching.

Finally, the parameters from the small PCL-only microcapsules (*t*_*c*_, *ε*, and *τ*_*f*_ ), the scaling factor *sf* from the small chitosan–PCL systems, and the mass transfer rate *k*_*c,P*_ from the large PCL-only microcapsules were initially applied to the large chitosan–PCL systems. Unlike the underprediction observed in large PCL-only capsules, the large chitosan-PCL microcapsules consistently overpredicted release, likely due to electrostatic interactions between chitosan and BSA, which slows diffusion. To account for this effect, *k*_*c,C*_ was estimated while keeping *t*_*c*_ = 28 days fixed. The last row in Figure 3 shows that overall release was well captured, with the exception of the 10% salt leaching formulation. This is likely due to the higher scaling factor *sf* used for this formulation. Furthermore, there was no clear trend on estimated *k*_*c,C*_ with increases in porosity (Table 3).

For bevacizumab, parameter estimation followed the same approach as for BSA. As shown in the first row of Figure 4, the model reproduced the experimental release across all PCL-only formulations. The estimated parameters (Table 4) revealed consistent trends, with higher salt leaching concentrations yielding increased porosity and reduced tortuosity.

Bevacizumab release from small chitosan-PCL microcapsules agreed well with the experimental data for the 5% and 7.5% formulations; however, the 10% salt leaching formulation missed some of the earlier and later time points (Figure 4, second row). The 7.5% and 10% formulations yielded similar scaling factors (Table 4), confirming the consistent delaying effect of chitosan. In contrast, the 5% formulation showed an *sf* near unity, suggesting a release profile similar to PCL-only. The corresponding confidence interval extended slightly below unity, reflecting the estimated weak influence of the chitosan layer. As shown in the bottom right panel in Figure B2, this near-unity scaling factor aligns with the substantial overlap between the yellow release data. The effective diffusivities in chitosan are expected to be similar among formulations, given that the sintering and salt leaching steps only affect PCL. Chitosan diffusivity in the first release phase 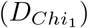 was close for the three formulations. Chitosan diffusivity during the second release phase 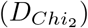 was similar for the 7.5 and 10% salt leaching formulations, but higher for the 5% formulation.

For the large PCL-only microcapsules, the initial validation without *k*_*c,P*_ and extended *t*_*c*_ also underpredicted bevacizumab release (Figure E12). After incorporating partial release from the capsule caps and setting *t*_*c*_ = 28 days, the model reproduced the experimental behavior well (Figure 4, third row), though the 7.5% and 10% formulations missed one or two data points. The estimated mass transfer rates *k*_*c,P*_ (Table 4) followed the expected trend, with higher salt concentrations yielding larger values consistent with enhanced release.

Bevacizumab release from the large chitosan-PCL microcapsules matched the experimental data for the 7.5% and 10% salt leaching formulations, while the predictions for the 5% formulation missed some of the data points. This behavior aligns with the lower scaling factor previously estimated for the 5% small chitosan–PCL capsules, which likely reduced the delaying effect of chitosan. As expected, higher salt concentrations corresponded to larger *k*_*c,C*_ values (Table 4), consistent with enhanced release from more porous devices.

### 3.2 Design exploration

For PCL-only microcapsules at a fixed total radius of 0.23 mm, thickening of the PCL layer while reducing the core radius consistently decreased both the therapeutic release rate duration and the cumulative drug release (Figures 5 and 6). The longest therapeutic release, lasting up to about 205 days, was achieved with the thinnest PCL layer (*δ*_*PCL*_ = 0.01 mm) and the largest core radius (*R*_*core*_ = 0.22 mm) tested in the 5% salt leaching formulation. In contrast, the 7.5% and 10% salt leaching formulations under the same geometry sustained therapeutic release for approximately 65 and 53 days, respectively. When the PCL thickness was increased to *δ*_*PCL*_ = 0.03 mm, cumulative release decreased due to the greater barrier imposed by the polymer layer. Under these conditions, the longest therapeutic release was observed for the highest salt leaching concentration, with the 10% formulation maintaining release for up to 125 days. Further increases in PCL thickness only shortened the duration of therapeutic release, as the larger polymer barrier reduced cumulative drug release. At the maximum thickness explored (*δ*_*PCL*_ = 0.09 mm), none of the formulations sustained therapeutic release rates.

**Fig. 5.**
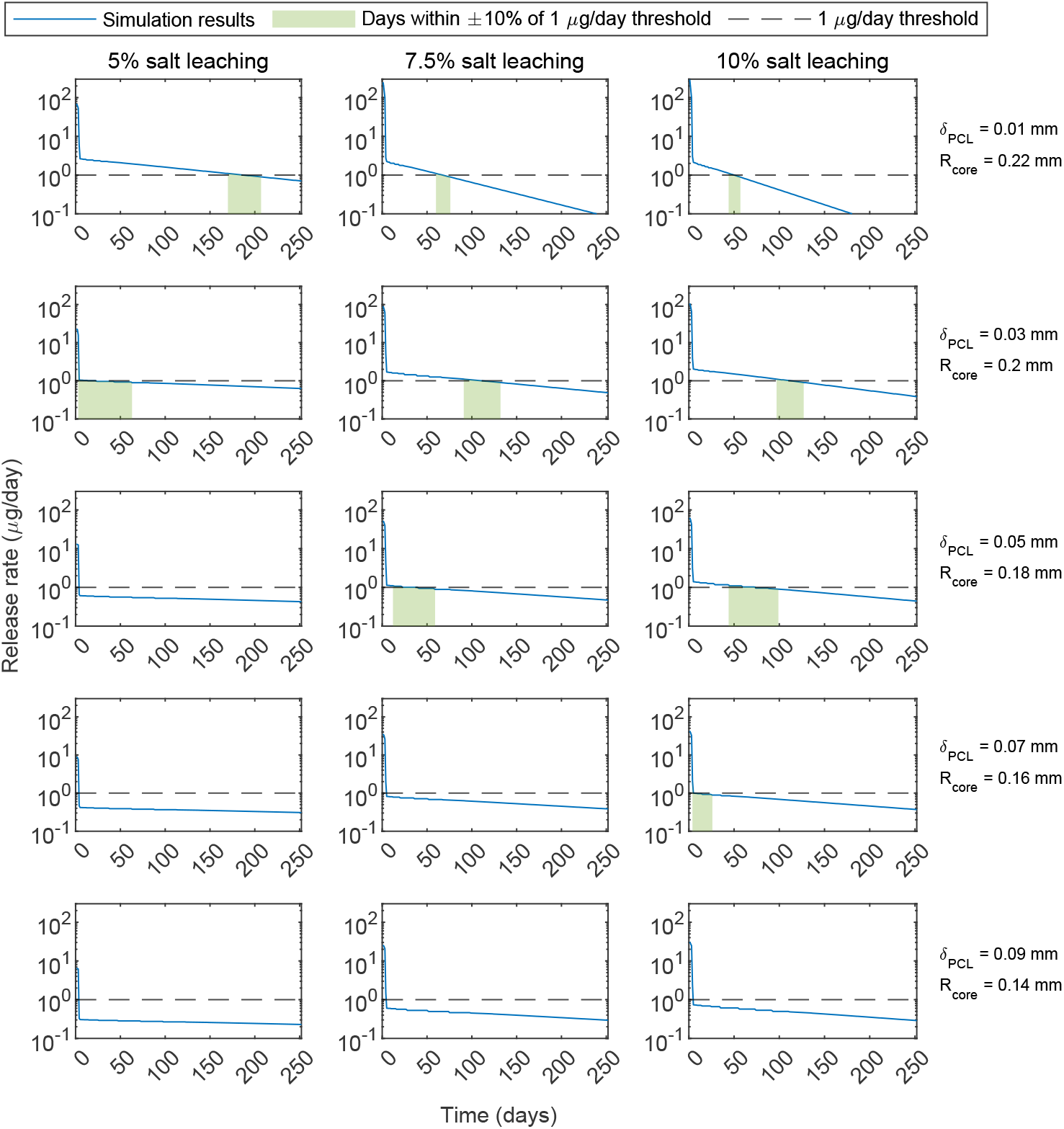
Bevacizumab release rate from different PCL-only designs with varying *δ*_*PCL*_ and core radius *R*_*core*_. Each panel shows profiles for different PCL-only configurations for a fixed 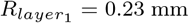. 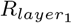 PCL radius. PCL: polycaprolactone. *δ*: thickness.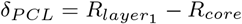.

**Fig. 6.**
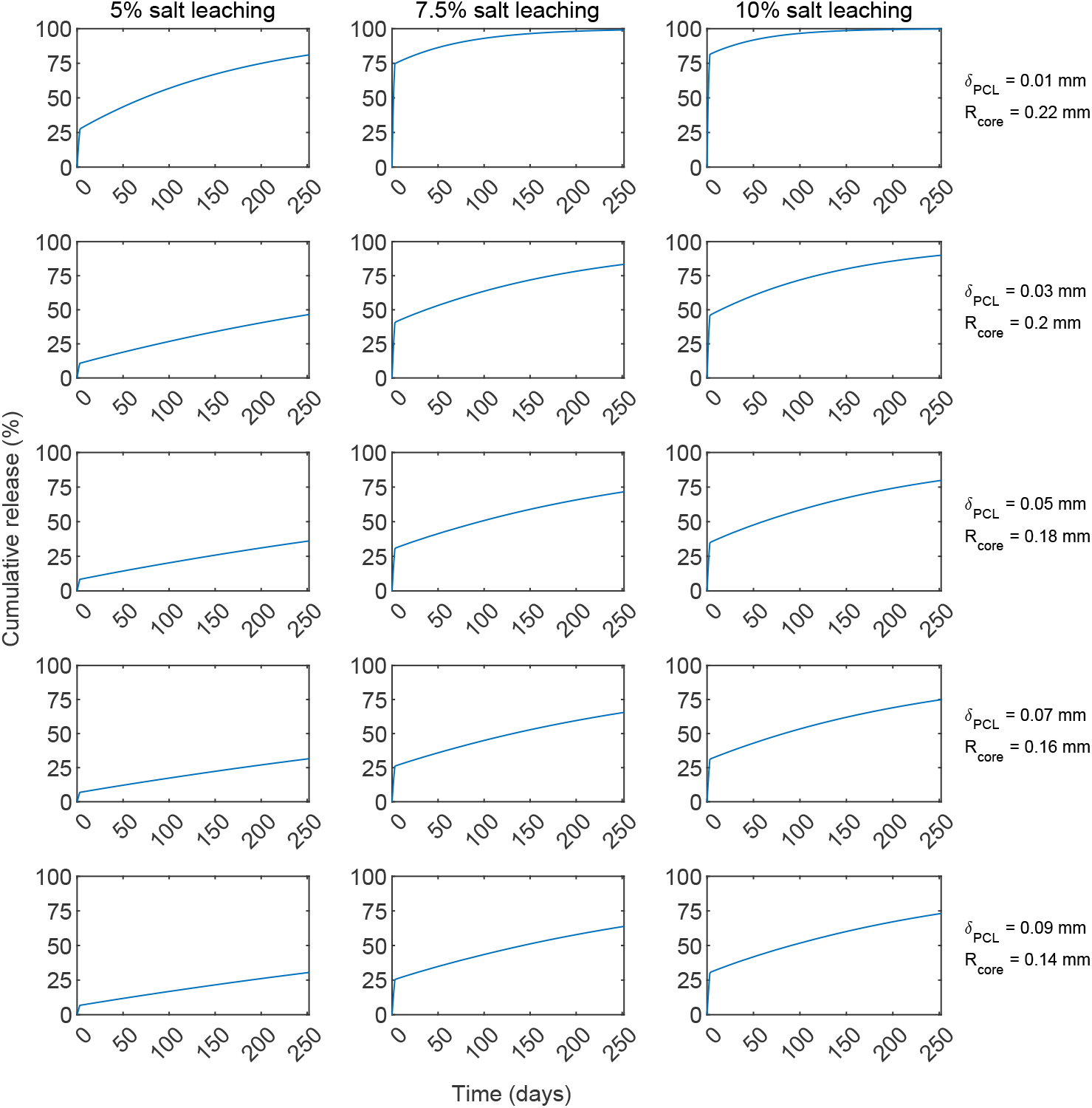
Cumulative bevacizumab release from different PCL-only designs with varying *δ*_*PCL*_ and core radius *R*_*core*_. Each panel shows profiles for different PCL-only configurations for a fixed 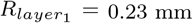. 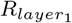 PCL radius. PCL: polycaprolactone. *δ*: thickness. 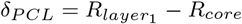.

In the case of chitosan–PCL microcapsules with a fixed composition of 75% PCL and 25% chitosan (*δ*_*PCL*_ : *δ*_*Chi*_ = 7.5 : 2.5), increasing the overall polymeric thickness at the expense of the core radius led to shorter therapeutic release durations and lower cumulative release values (Figures 7 and 8). At the smallest thickness tested (*δ*_*PCL*_ + *δ*_*Chi*_ = 0.01 mm), the 5% salt leaching formulation achieved the longest therapeutic release, extending to approximately 208 days, whereas the 7.5% and 10% formulations sustained release for about 145 and 134 days, respectively. When the total thickness was raised to *δ*_*PCL*_ + *δ*_*Chi*_ = 0.03 mm, therapeutic performance declined, with the 10% salt leaching formulation maintaining release for only around 125 days. For polymeric layers of *δ*_*PCL*_ + *δ*_*Chi*_ ≥ 0.05 mm, drug release rates fell below the therapeutic release rate threshold, despite more than half of the drug remaining in the DDSs after 250 days.

**Fig. 7.**
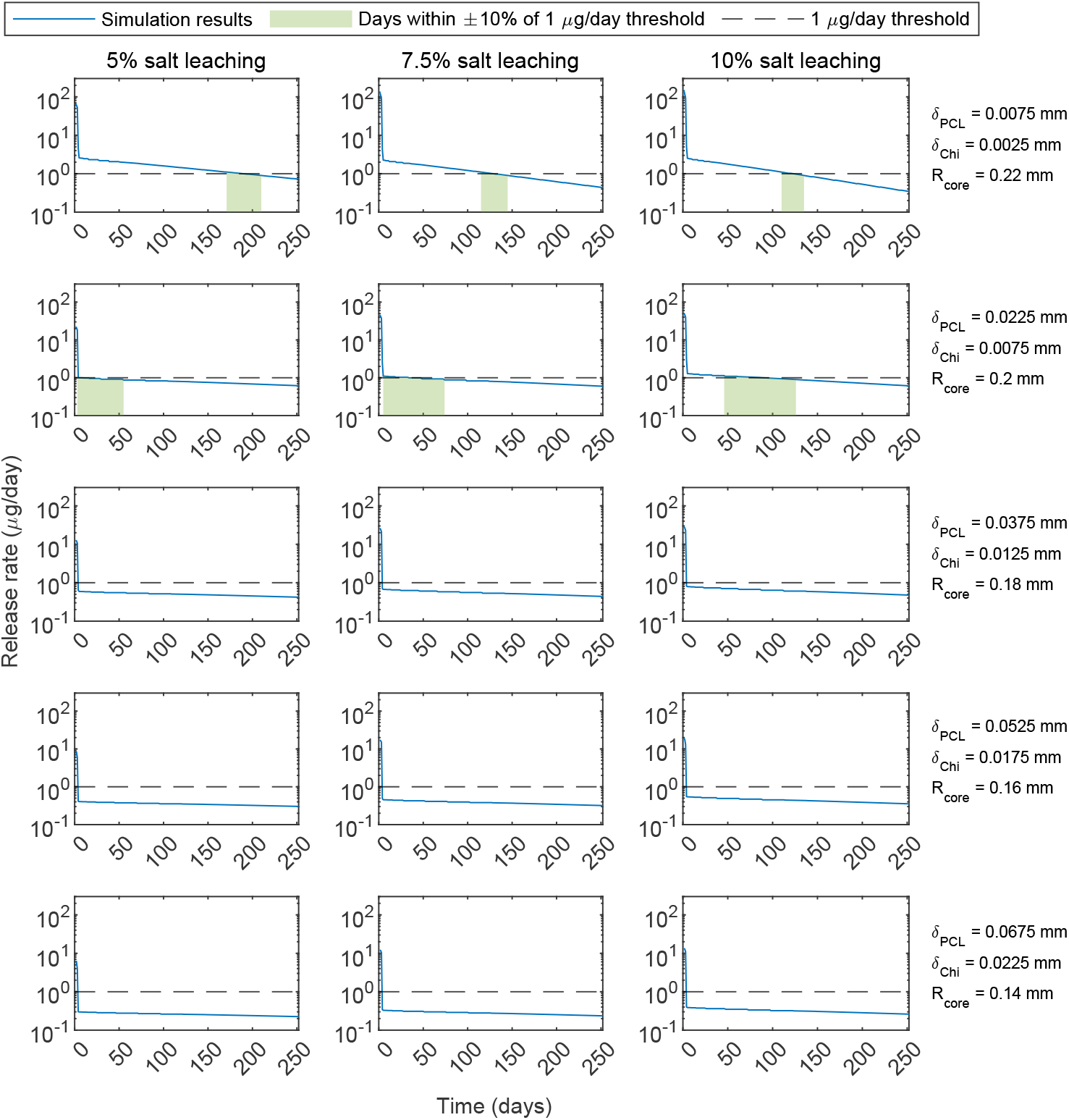
Bevacizumab release rate from different chitosan-PCL designs with a fixed composition of 75% PCL and 25% chitosan (*δ*_*PCL*_ : *δ*_*Chi*_ = 0.75 : 0.25) and varying total polymer thickness *δ*_*PCL*_ + *δ*_*Chi*_ and core radius *R*_*core*_. Each panel shows profiles for different configurations for a fixed 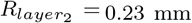. 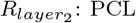 PCL radius. 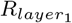 chitosan radius. PCL: polycaprolactone. Chi: chitosan. *δ*: thickness. 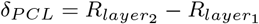 and 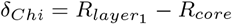.

**Fig. 8.**
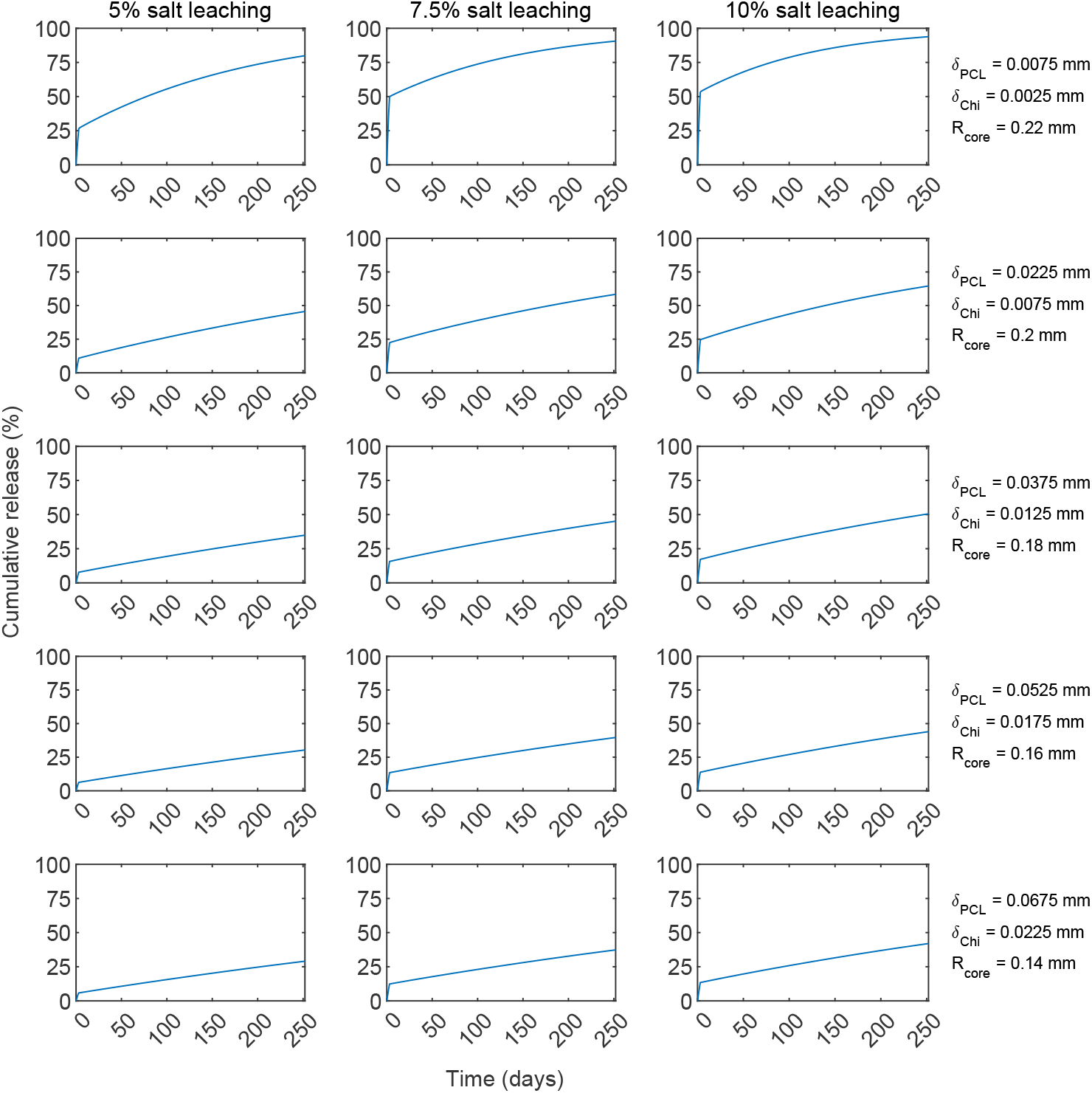
Cumulative bevacizumab release from different chitosan-PCL designs with a fixed composition of 75% PCL and 25% chitosan (*δ*_*PCL*_ : *δ*_*Chi*_ = 0.75 : 0.25) and varying total polymer thickness *δ*_*PCL*_ + *δ*_*Chi*_ and core radius *R*_*core*_. Each panel shows profiles for different configurations for a fixed 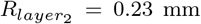. 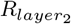 PCL radius. 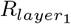 chitosan radius. PCL: polycaprolactone. Chi: chitosan. *δ*: thickness. 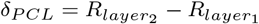 and 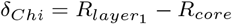.

For chitosan–PCL microcapsules with a fixed total polymeric thickness of *δ*_*PCL*_ + *δ*_*Chi*_ = 0.01 mm, reducing the proportion of PCL while increasing the chitosan fraction extended the duration of therapeutic bevacizumab release, though at the expense of cumulative release (Figures 9 and 10). This outcome reflects the lower effective diffusivity of chitosan compared to PCL, which slows drug transport through the capsule wall. At the most chitosan-rich composition tested (*δ*_*PCL*_ : *δ*_*Chi*_ = 1 : 9), therapeutic release persisted for approximately 220 days with 5% salt leaching, nearly 200 days with 10%, and about 175 days with 7.5%. The counterintuitive ordering between the 7.5% and 10% formulations can be explained by the estimated scaling factors, which were lower for the 10% formulation (*sf* = 5.62) compared to the 7.5% formulation (*sf* = 4.89), thereby slowing drug release and extending therapeutic duration. Increasing the relative proportion of PCL consistently shortened therapeutic release across all formulations. Nevertheless, at the fixed thickness of *δ*_*PCL*_ +*δ*_*Chi*_ = 0.01 mm, every formulation maintained therapeutic levels for at least 90 days, with the 5% salt leaching condition emerging as the most favorable, regardless of the PCL:chitosan ratio.

**Fig. 9.**
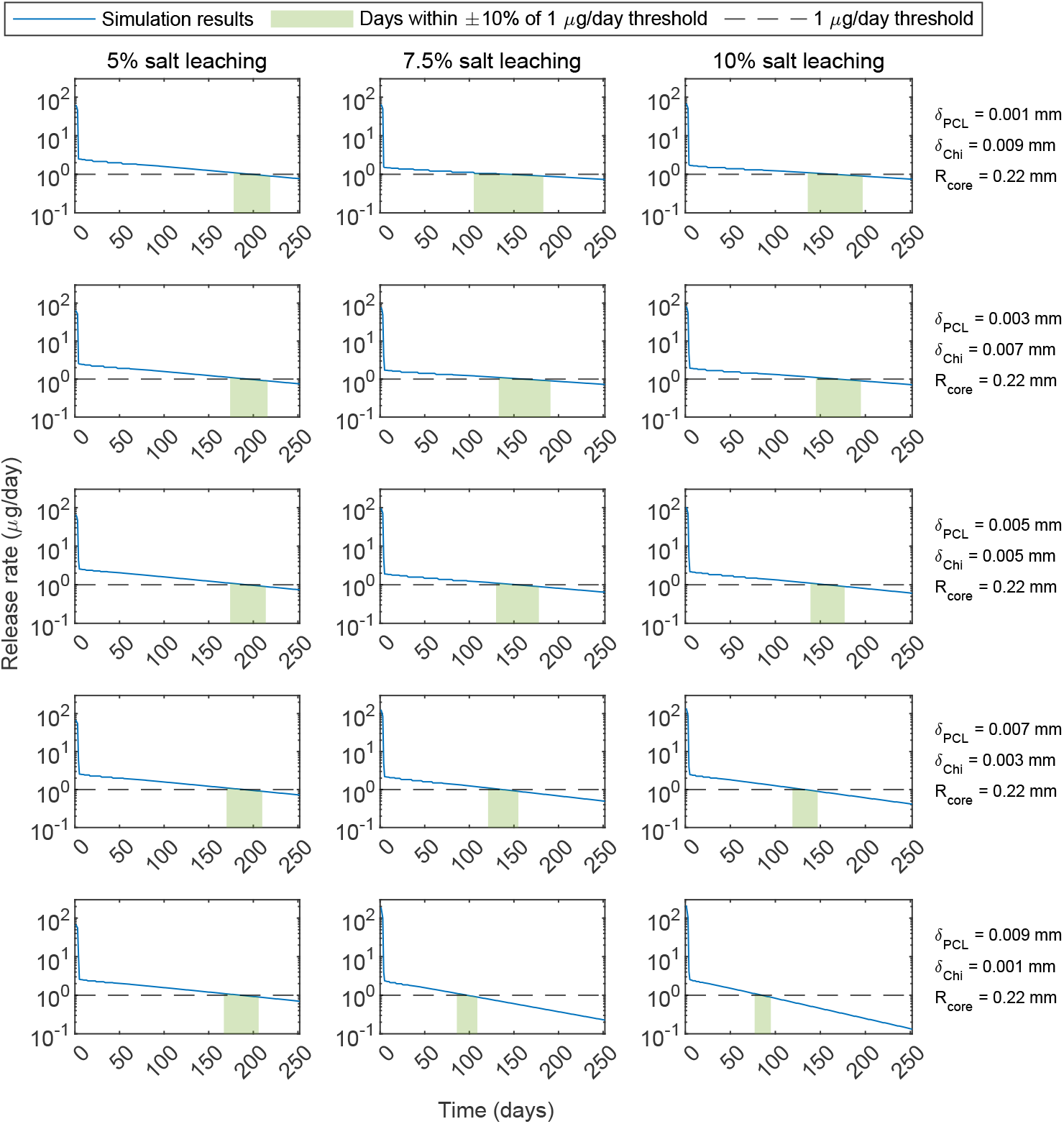
Bevacizumab release rate from different chitosan-PCL designs with a fixed polymer thickness of *δ*_*PCL*_ + *δ*_*Chi*_ = 0.01 mm and core radius *R*_*core*_ = 0.22 mm and varying PCL:chitosan ratio *δ*_*PCL*_ : *δ*_*Chi*_. Each panel shows profiles for different PCL-only configurations for a fixed 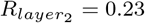 mm. 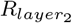 PCL radius. 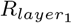 chitosan radius. PCL: polycaprolactone. Chi: chitosan. *δ*: thickness. 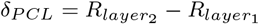 and 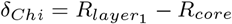.

**Fig. 10.**
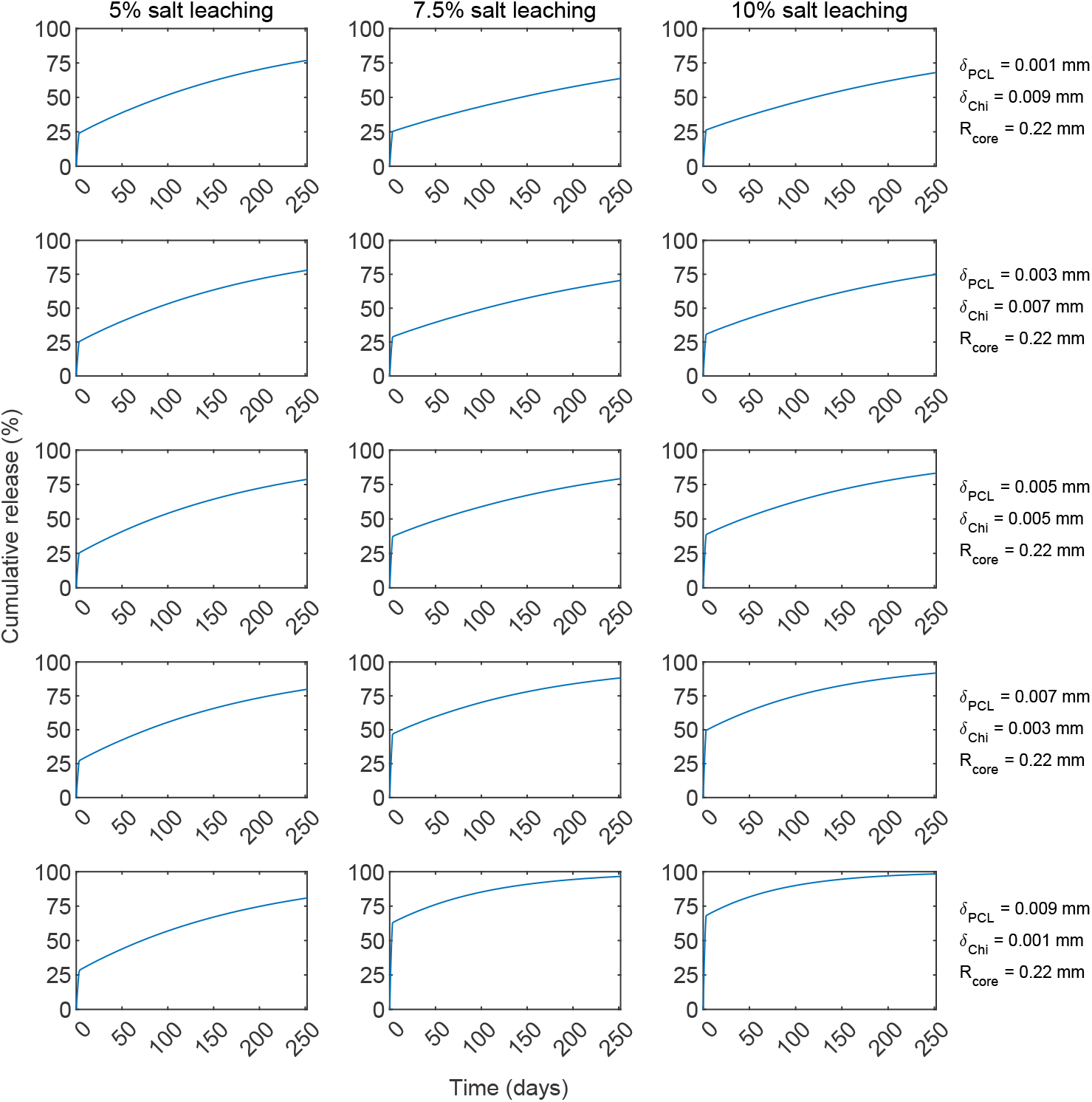
Cumulative bevacizumab release from different chitosan-PCL designs with fixed polymer thickness of *δ*_*PCL*_ + *δ*_*Chi*_ = 0.01 mm and core radius *R*_*core*_ = 0.22 mm and varying PCL:chitosan ratio *δ*_*PCL*_ : *δ*_*Chi*_. Each panel shows profiles for different chitosan-PCL configurations for a fixed 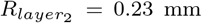. 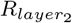 PCL radius. 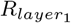 chitosan radius. PCL: polycaprolactone. Chi: chitosan. *δ*: thickness. 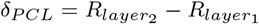 and 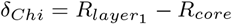.

## 4 Discussion

This study established a computational framework that integrates diffusion-controlled modeling, progressive parameter estimation, and uncertainty quantification across microcapsule designs and formulations of PCL-only and chitosan-PCL microcapsules. By anchoring the parametrization to the simplest system (small PCL-only microcapsules) and progressively extending to more complex designs, the framework captured both baseline release behavior and the additional effects introduced by the chitosan layer and imperfect caps sealing.

Salt leaching concentration played a central role in shaping porosity and tortuosity, with higher concentrations increasing diffusivity and enhancing release. However, variability in pore formation introduced deviations, where in some cases initial release was faster for lower salt leaching concentrations or the total release for the maximum salt leaching concentration was below the expected when compared to the other two formulations, disrupting the expected trends if the release were monotonically dependent on salt leaching concentration. This experimental variability is consistent with reported similarity in pore size distributions [11] (Figure A1).

Comparing the two drugs highlighted distinct release dynamics across formulations. Bevacizumab exhibited faster initial release (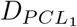 and 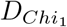) than BSA in the 7.5% and 10% salt leaching formulations, whereas BSA release was faster during the second phase (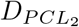 and 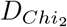). These behaviors reflect differences in drug molecular size and interactions with the chitosan matrix, consistent with prior observations [27]. The scaling factors further emphasized this distinction: values were generally higher for BSA, underscoring the stronger delaying effect of chitosan on its diffusion compared to bevacizumab. Interestingly, the 5% chitosan–PCL formulation for bevacizumab produced a scaling factor around unity, suggesting that the chitosan influence was limited in this particular formulation.

Model performance also differed between drugs and formulations. In general, bevacizumab release profiles aligned more closely with experimental data, yielding lower RMSE values than BSA in most cases. For large capsules, incorporating mass transfer rates *k*_*c,i*_ was essential to account for partial release through the caps. In PCL-only systems, higher salt concentrations corresponded to larger values, consistent with enhanced release from more porous devices. In contrast, large chitosan–PCL capsules required estimation of *k*_*c,C*_ to account for electrostatic interactions between chitosan and BSA, which slowed diffusion and led to overprediction when baseline parameters were applied directly.

Design variations further clarified the dominant role of both total polymer thickness and PCL:chitosan ratio in controlling release behavior. In PCL-only microcapsules, reducing the PCL thickness while enlarging the core radius consistently increased cumulative release and, depending on the formulation, could also extend therapeutic release duration. Importantly, the effect of thickness was not independent as the salt leaching concentration also played a role in drug release behavior and both need to be considered together. In chitosan–PCL DDSs, thinning the total polymeric layer at a fixed PCL:chitosan ratio consistently prolonged therapeutic release across all formulations. At the smallest total polymeric layer examined, decreasing the PCL:chitosan ratio further enhanced the duration of release within the therapeutic threshold, reflecting the slower diffusivity of chitosan relative to PCL. These outcomes are closely tied to the estimated scaling factors: the 5% salt leaching formulation yielded a value near unity, whereas the 7.5% and 10% formulations were closer to five. This distinction is critical, as approximating the 5% *sf* to higher values would imply slower release kinetics and potentially shorter therapeutic duration than observed.

Although the results presented here are encouraging, experimental validation remains a critical next step. One key assumption in the design exploration is that polymer thickness does not alter effective diffusivity. In practice, however, changes in thickness also affect tortuosity, which may lead to release profiles that differ from those predicted. To address this, new experiments should be conducted using one of the proposed designs, with release monitored over time. While the current analysis focused on cumulative release over 250 days, validation could be performed over shorter intervals (30–60 days), allowing direct comparison between simulated and experimental profiles. Such studies would either confirm the model’s predictive capability or provide new data to refine parameterization when thickness effects are incorporated.

Another limitation is that the present work relies on *in vitro* release data, whereas *in vivo* conditions can accelerate polymer degradation and alter release kinetics [17, 36]. Furthermore, the therapeutic threshold in this study was defined in terms of release rate, but a more clinically relevant metric is the drug concentration at the macula. Future work could, therefore, integrate this release model with our three-dimensional ocular pharmacokinetics framework for rabbit and human eyes [37], enabling direct prediction of macular concentration.

## 5 Conclusion

This work established a computational framework that integrates diffusion-driven models, progressive parameter estimation, and uncertainty quantification to describe drug release from multi-layered microcapsules. Starting from the simplest microcapsule design and progressively extending to more complex systems, the framework captured baseline release behavior and any additional effects introduced in more complex designs. Salt leaching level was identified as a key driver of porosity and diffusivity; however, variability in pore formation occasionally disrupted expected trends. Distinct release patterns between BSA and bevacizumab highlighted the influence of drug size and drug interactions with chitosan, with scaling factors suggesting stronger delaying effects of chitosan on BSA. Incorporating mass transfer rates was essential for reproducing release in larger particles, and the magnitude was different depending on the presence of chitosan, which slowed down release. Design exploration emphasized total polymer thickness and PCL:chitosan ratio as the dominant control variables, with increased core size providing greater load amount capability. The simultaneous reduction of total polymeric thickness and PCL:chitosan ratio in the chitosan-PCL microcapsules, enabled extended therapeutic release regardless of formulations, with the overall best design being the 5% salt leaching formulation with a total polymeric thickness of 0.01 mm and 1:9 PCL:chitosan ratio. This formulation provided longer therapeutic release and lower cumulative drug release than the analogous PCL-only microcapsule. While these findings demonstrate the promise of chitosan–PCL microcapsules in moderating burst release and supporting long-acting delivery, further validation is needed, including experiments with new designs. The mechanistic framework developed here can be easily adapted to other multi-layered microcapsule systems, and can be used to inform the design of long-lasting drug delivery systems.

## Supplementary information

If your article has accompanying supplementary file/s please state so here.

Please refer to Journal-level guidance for any specific requirements.

## Acknowledgements

This work was supported by National Institutes of Health grant R01EB032870 to KESR and ANFV, Owen Locke Foundation to KESR, and the University at Buffalo, an Owen Locke Foundation grant to KESR, an Ohio Lions Eye Research Foundation grant to KESR, The Ohio State University College of Engineering, and the University at Buffalo.

## Appendix A Pore diameter distribution in microcapsules

**Fig. A1.**
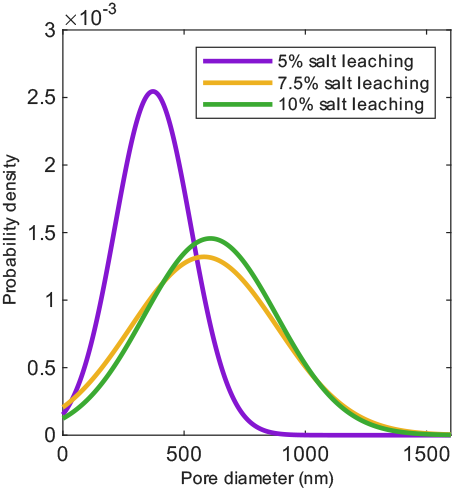
Normal distribution of pore diameters in the polycaprolactone layer after the microcapsule fabrication process. The reported average pore diameters ± standard deviations for the three pore sizes are: i) 5% salt leaching: 371.65 ± 156.77 nm, ii) 7.5% salt leaching: 582.21 ± 302.17 nm, and iii) 10% salt leaching: 608.55 ± 273.90 nm [11].

## Appendix B Experimental drug release data

**Fig. B2.**
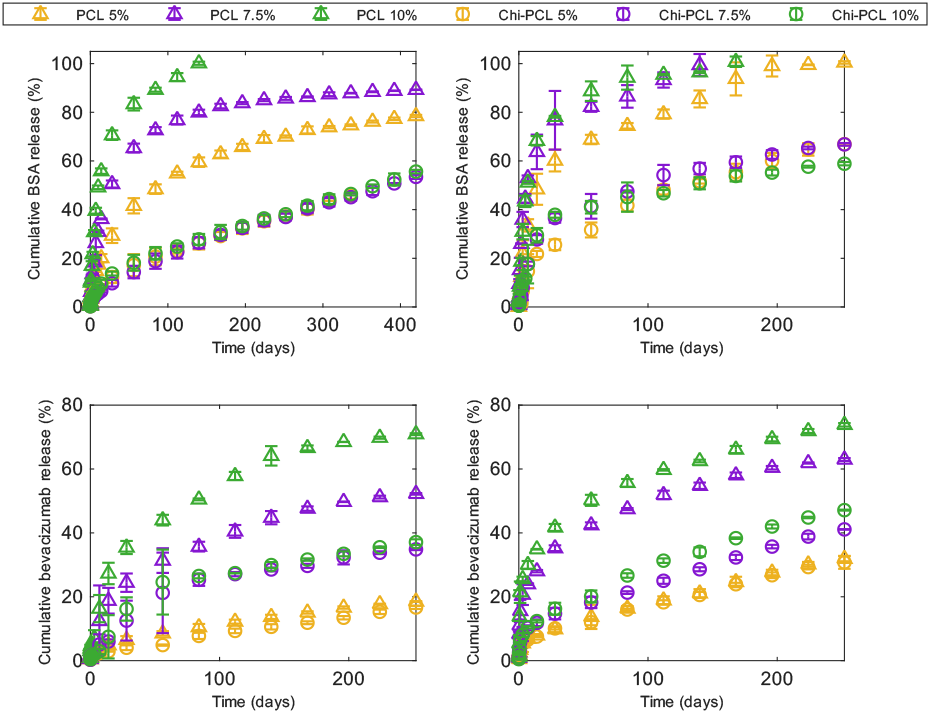
Experimental cumulative drug release profiles for different formulations and drugs from Jiang et al. [11]. Top row: bovine serum albumin release. Bottom row: bevacizumab release. First column: large microcapsules (*R*_*core*_ = 0.8225 mm). Second column: small microcapsules (*R*_*core*_ = 0.13 mm). Percentage refers to salt leaching percentage of HEPES salts added. *R*_*core*_: core radius. PCL: polycaprolactone. Chi: chitosan.

## Appendix C Preliminary parameter estimation on PCL-only small microcapsules

Figure C3 shows the cumulative BSA release profile for small microcapsules with 5% salt leaching evaluated at fixed critical times of *t*_*c*_ = 3, 7, and 14 days. The case of *t*_*c*_ = 3 days yields the highest error values, while the simulations with *t*_*c*_ = 7 and *t*_*c*_ = 14 days produce similar performance. Across all critical times, the average and best model predictions overlap and reproduce the overall shape of the experimental release curves. The agreement with the experimental data is particularly strong for the *t*_*c*_ = 7 and *t*_*c*_ = 14 days. Table C1 highlights the close correspondence between the parameters obtained from the best and average models.

**Fig. C3.**
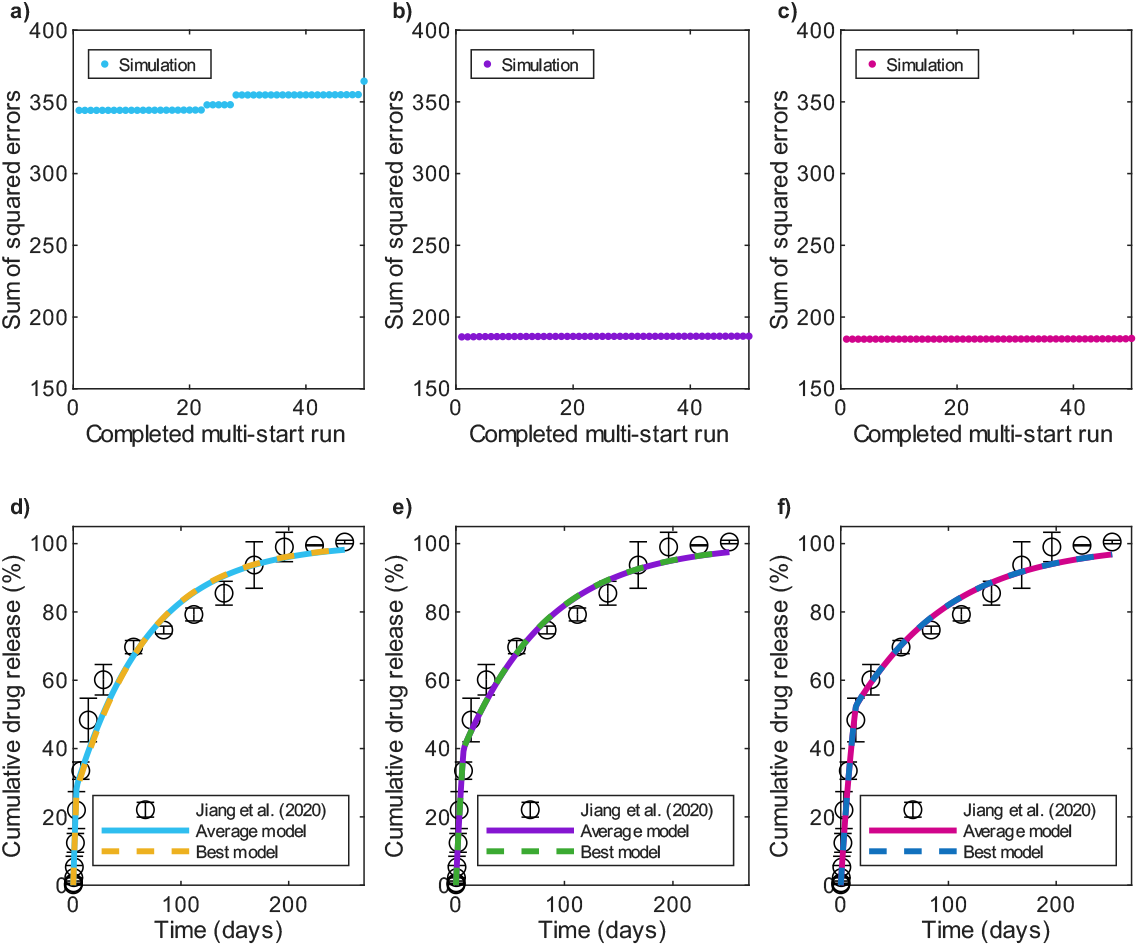
Cumulative BSA release profiles after multi-start parameter estimation in 5% salt leaching PCL-only microcapsules with parameters in Table C1. First column: *t*_*c*_ = 3 days. Second column: *t*_*c*_ = 7 days. Third column: *t*_*c*_ = 14 days. First row: Error values. Second row: Average and best models compared to experimental data. Experimental data from Jiang et al. [11]. BSA: bovine serum albumin. PCL: Polycaprolactone.

Figure C4 presents the cumulative BSA release profiles for small microcapsules with 7.5% salt leaching evaluated at fixed critical times of *t*_*c*_ = 3, 7, and 14 days. In this formulation, larger critical times correspond to higher error values (Table C1), indicating diminished model performance as *t*_*c*_ increases. While across all scenarios the average and best models overlap and are comparable to the experimental data, the case of *t*_*c*_ = 3 days is predominantly better performing.

**Fig. C4.**
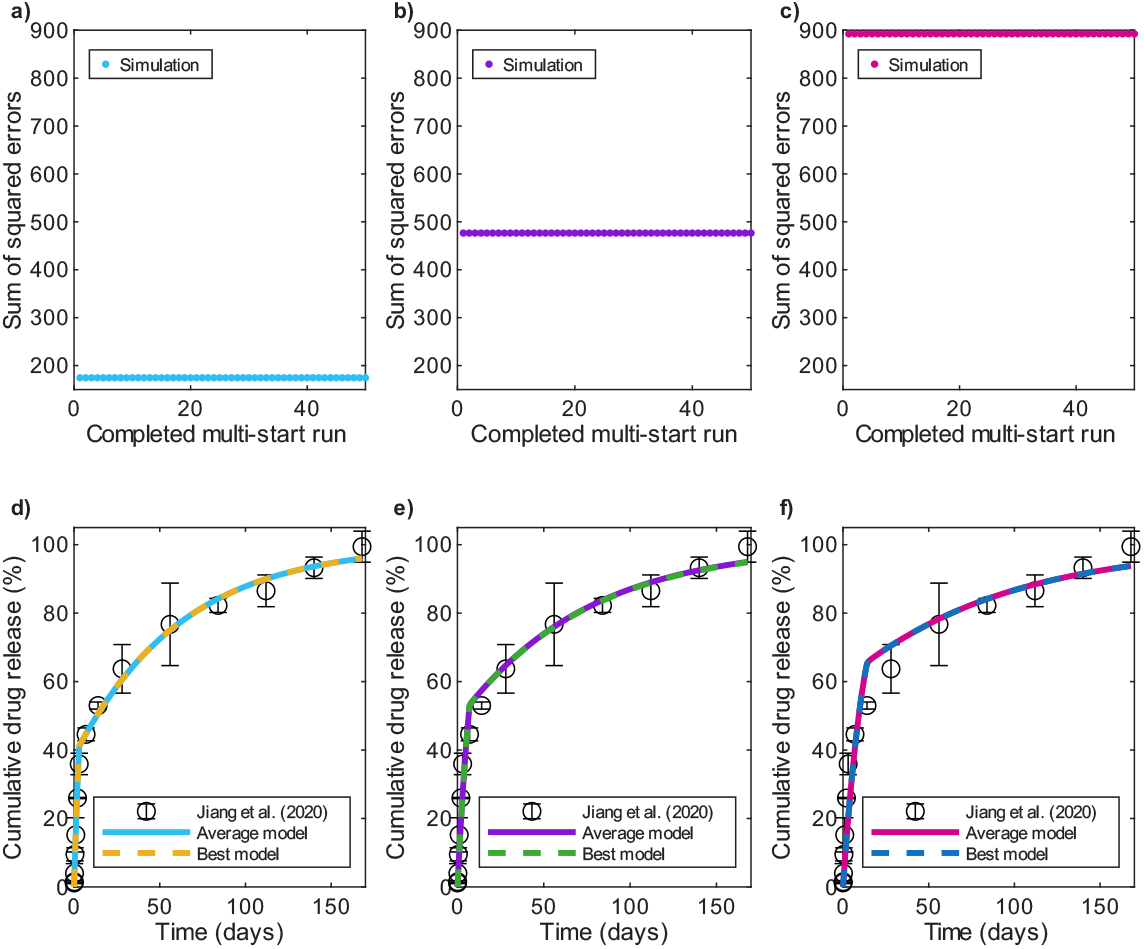
Cumulative BSA release profiles after multi-start parameter estimation in 7.5% salt leaching PCL-only microcapsules with parameters in Table C1. First column: *t*_*c*_ = 3 days. Second column: *t*_*c*_ = 7 days. Third column: *t*_*c*_ = 14 days. First row: Error values. Second row: Average and best models compared to experimental data. Experimental data from Jiang et al. [11]. BSA: bovine serum albumin. PCL: Polycaprolactone.

Figure C5 shows the cumulative BSA release profiles for small microcapsules with 10% salt leaching evaluated at fixed critical times of *t*_*c*_ = 3, 7, and 14 days. Consistent with the 5% salt leaching formulation, the case with *t*_*c*_ = 3 days shows the largest deviation from the experimental data. In contrast, the simulations with *t*_*c*_ = 7 and *t*_*c*_ = 14 days yield similar performance, with *t*_*c*_ = 7 days case providing slightly better agreement to the data. There is no appreciable difference between average models and best models at the different critical times selected, and the parameter estimates obtained from both models are closely aligned (Table C1).

Figure C6 presents the cumulative bevacizumab release profile for small microcapsules with 5% salt leaching evaluated at fixed critical times of *t*_*c*_ = 3, 7, and 14 days. In this formulation, larger critical times correspond to higher error values, although the differences remain relatively small. Among the different cases, *t*_*c*_ = 3 days yields the lowest error. In all cases the parameters from average and best models align closely (Table C1), resulting in successful reproduction of the linear release behavior observed in the experimental data.

**Fig. C5.**
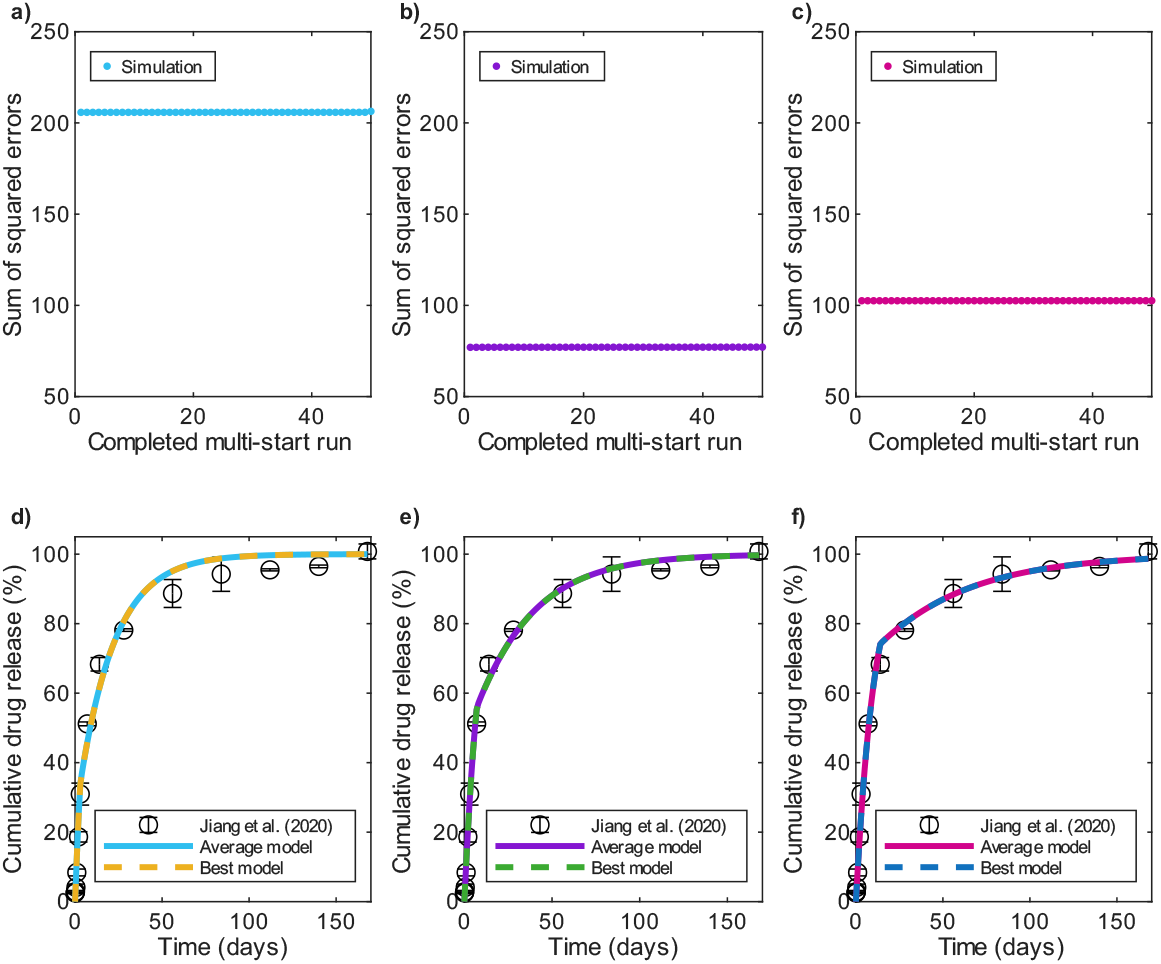
Cumulative BSA release profiles after multi-start parameter estimation in 10% salt leaching PCL-only microcapsules with parameters in Table C1. First column: *t*_*c*_ = 3 days. Second column: *t*_*c*_ = 7 days. Third column: *t*_*c*_ = 14 days. First row: Error values. Second row: Average and best models compared to experimental data. Experimental data from Jiang et al. [11]. BSA: bovine serum albumin. PCL: Polycaprolactone.

**Fig. C6.**
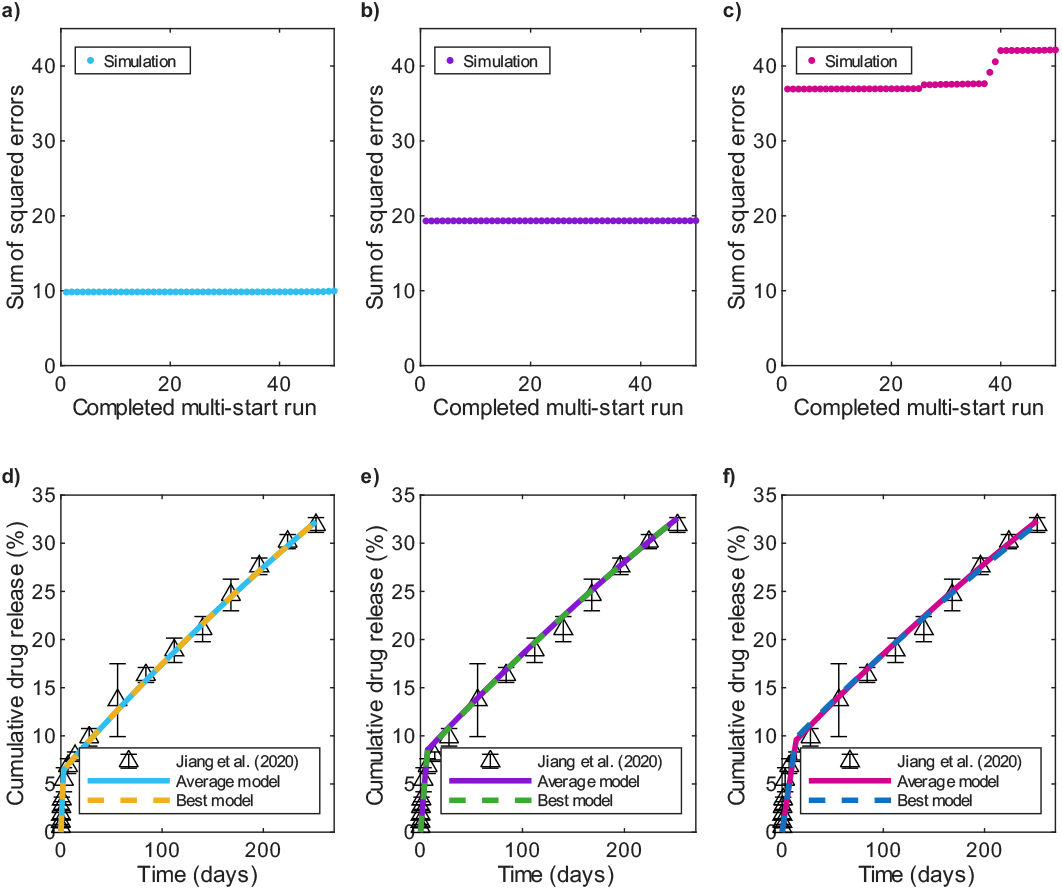
Cumulative bevacizumab release profiles after multi-start parameter estimation in 5% salt leaching PCL-only microcapsules with parameters in Table C1. First column: *t*_*c*_ = 3 days. Second column: *t*_*c*_ = 7 days. Third column: *t*_*c*_ = 14 days. First row: Error values. Second row: Average and best models compared to experimental data. Experimental data from Jiang et al. [11].

Figure C7 shows cumulative bevacizumab release profiles for small microcapsules with 7.5% salt leaching evaluated at fixed critical times of *t*_*c*_ = 3, 7, and 14 days. As in the 5% salt leaching formulation, the lowest error values correspond to the lowest critical times; however, in this case the differences between errors are more pronounced. For *t*_*c*_ = 3 and 14 days, the average and best model predictions coincide, whereas at *t*_*c*_ = 7 days a slight divergence is observed between the two. Overall, *t*_*c*_ = 3 days case provides the best agreement with the experimental data (Table C1).

**Fig. C7.**
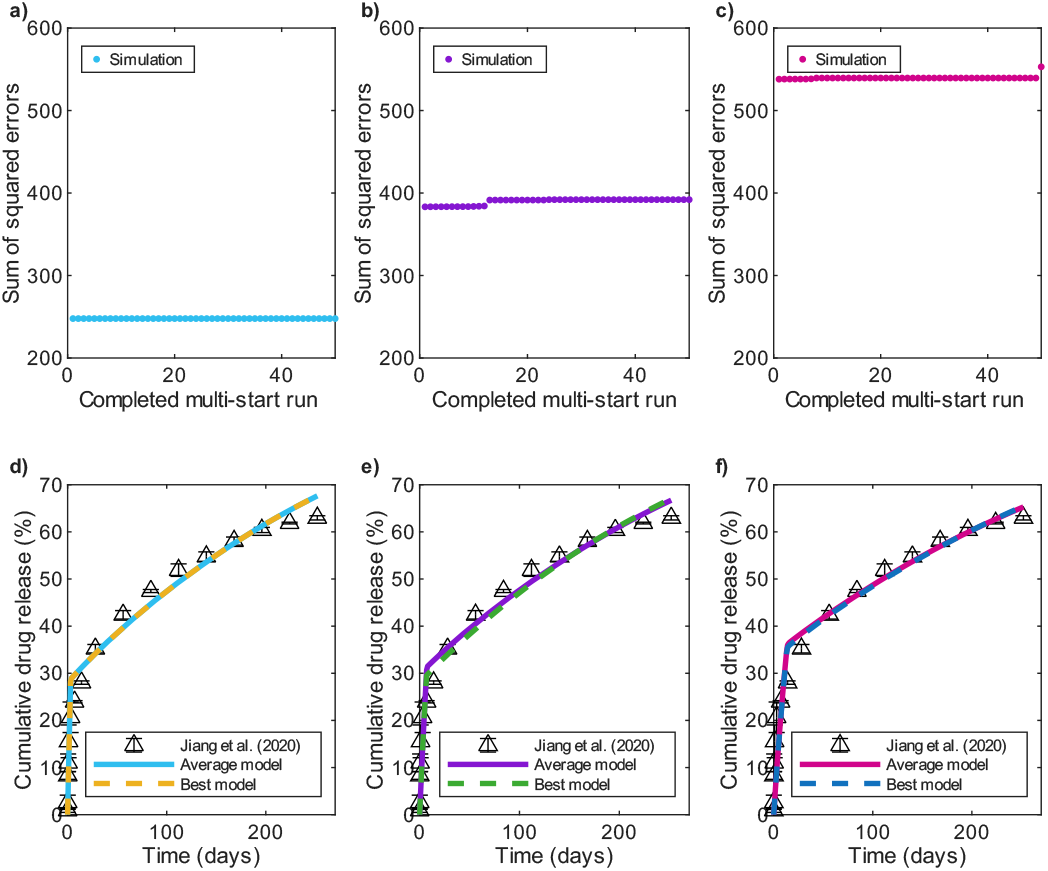
Cumulative bevacizumab release profiles after multi-start parameter estimation in 7.5% salt leaching PCL-only microcapsules with parameters in Table C1. First column: *t*_*c*_ = 3 days. Second column: *t*_*c*_ = 7 days. Third column: *t*_*c*_ = 14 days. First row: Error values. Second row: Average and best models compared to experimental data. Experimental data from Jiang et al. [11].

Figure C8 shows cumulative bevacizumab release profiles for small microcapsules with 10% salt leaching evaluated at fixed critical times of *t*_*c*_ = 3, 7, and 14 days. Consistent with the trend observed in the other bevacizumab formulations, the *t*_*c*_ = 3 days case yields the lowest error values while *t*_*c*_ = 14 days produces the largest errors. For *t*_*c*_ = 3 days, the average and best model predictions coincide, while at *t*_*c*_ = 7 and 14 days slight deviations are evident between average and best model predictions. Although all cases capture the overall shape of the experimental release curve, the *t*_*c*_ = 3 days case provides the closest agreement to the data (Table C1).

**Table C1.** Estimated parameters for fitting the model to data for small drug-loaded PCL-only microcapsules from Jiang et al. [35] at different critical times. The average model is obtained by averaging the parameters obtained from all the optimization runs that achieved an error within 5% of the minimum error after 50 multi-start attempts. The best model corresponds to the parameter set with the lowest error value. The critical time *t*_*c*_ is fixed at the different values shown. BSA: bovine serum albumin. *t*_*c*_: critical time for phase transition. *ε*: PCL porosity. 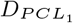 drug diffusivity in PCL during the first release phase. *τ*_2_: PCL tortuosity during the second release phase. 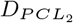 drug diffusivity in PCL during the second release phase. S. L.: salt leaching. ^∗^: calculated values.

| BSA: best results |  |  |  |  |  |  |
| --- | --- | --- | --- | --- | --- | --- |
| S. L. (%) | $t_c$ (days) | $\epsilon$ | $D_{PCL_1}$ (cm<sup>2</sup>/s)* | $\tau_2$ | $D_{PCL_2}$ (cm<sup>2</sup>/s)* | Error value |
| 5 | 3 | $8.15 \times 10^{-3}$ | $5.59 \times 10^{-11}$ | 894.6 | $7.66 \times 10^{-12}$ | 344.1 |
| | 7 | $6.87 \times 10^{-3}$ | $3.97 \times 10^{-11}$ | 875.7 | $6.60 \times 10^{-12}$ | 191.0 |
| | 14 | $5.73 \times 10^{-3}$ | $2.76 \times 10^{-11}$ | 825.6 | $5.84 \times 10^{-12}$ | 184.6 |
| 7.5 | 3 | $1.07 \times 10^{-2}$ | $9.63 \times 10^{-11}$ | 1083 | $8.31 \times 10^{-12}$ | 174.5 |
| | 7 | $8.14 \times 10^{-3}$ | $5.57 \times 10^{-11}$ | 969.2 | $7.06 \times 10^{-12}$ | 476.6 |
| | 14 | $6.87 \times 10^{-3}$ | $3.97 \times 10^{-11}$ | 1009 | $5.73 \times 10^{-12}$ | 892.4 |
| 10 | 3 | $9.39 \times 10^{-3}$ | $7.42 \times 10^{-11}$ | 314.2 | $2.51 \times 10^{-11}$ | 205.8 |
| | 7 | $8.40 \times 10^{-3}$ | $5.93 \times 10^{-11}$ | 437.5 | $1.61 \times 10^{-11}$ | 80.00 |
| | 14 | $7.68 \times 10^{-3}$ | $4.96 \times 10^{-11}$ | 657.0 | $9.83 \times 10^{-12}$ | 102.5 |
| BSA: average results |  |  |  |  |  |  |
| S. L. (%) | $t_c$ (days) | $\epsilon$ | $D_{PCL_1}$ (cm<sup>2</sup>/s)* | $\tau_2$ | $D_{PCL_2}$ (cm<sup>2</sup>/s)* | Error value |
| 5 | 3 | $8.39 \times 10^{-3}$ | $5.92 \times 10^{-11}$ | 937.9 | $7.52 \times 10^{-12}$ | 349.4 |
| | 7 | $6.88 \times 10^{-3}$ | $3.98 \times 10^{-11}$ | 875.7 | $6.61 \times 10^{-12}$ | 191.0 |
| | 14 | $5.72 \times 10^{-3}$ | $2.75 \times 10^{-11}$ | 827.2 | $5.82 \times 10^{-12}$ | 184.7 |
| 7.5 | 3 | $1.07 \times 10^{-2}$ | $9.63 \times 10^{-11}$ | 1082 | $8.32 \times 10^{-12}$ | 174.6 |
| | 7 | $8.14 \times 10^{-3}$ | $5.57 \times 10^{-11}$ | 968.5 | $7.07 \times 10^{-12}$ | 476.6 |
| | 14 | $6.88 \times 10^{-3}$ | $3.98 \times 10^{-11}$ | 1009 | $5.73 \times 10^{-12}$ | 892.4 |
| 10 | 3 | $9.39 \times 10^{-3}$ | $7.42 \times 10^{-11}$ | 314.1 | $2.51 \times 10^{-11}$ | 205.8 |
| | 7 | $8.44 \times 10^{-3}$ | $5.99 \times 10^{-11}$ | 448.6 | $1.58 \times 10^{-11}$ | 80.11 |
| | 14 | $7.69 \times 10^{-3}$ | $4.97 \times 10^{-11}$ | 657.7 | $9.83 \times 10^{-12}$ | 102.5 |
| Bevacizumab: best results |  |  |  |  |  |  |
| S. L. (%) | $t_c$ (days) | $\epsilon$ | $D_{PCL_1}$ (cm<sup>2</sup>/s)* | $\tau_2$ | $D_{PCL_2}$ (cm<sup>2</sup>/s)* | Error value |
| 5 | 3 | $4.36 \times 10^{-3}$ | $1.36 \times 10^{-11}$ | 4658 | $6.70 \times 10^{-13}$ | 9.81 |
| | 7 | $3.06 \times 10^{-3}$ | $6.70 \times 10^{-12}$ | 3376 | $6.49 \times 10^{-13}$ | 19.32 |
| | 14 | $2.36 \times 10^{-3}$ | $3.99 \times 10^{-12}$ | 2739 | $6.17 \times 10^{-13}$ | 36.93 |
| 7.5 | 3 | $8.95 \times 10^{-3}$ | $5.74 \times 10^{-11}$ | 3865 | $1.66 \times 10^{-12}$ | 247.8 |
| | 7 | $6.16 \times 10^{-3}$ | $2.72 \times 10^{-11}$ | 2758 | $1.60 \times 10^{-12}$ | 383.3 |
| | 14 | $4.89 \times 10^{-3}$ | $1.71 \times 10^{-11}$ | 2564 | $1.37 \times 10^{-12}$ | 538.0 |
| 10 | 3 | $9.96 \times 10^{-3}$ | $7.10 \times 10^{-11}$ | 3281 | $2.17 \times 10^{-12}$ | 270.5 |
| | 7 | $7.24 \times 10^{-3}$ | $3.75 \times 10^{-11}$ | 2737 | $1.89 \times 10^{-12}$ | 508.8 |
| | 14 | $5.52 \times 10^{-3}$ | $2.18 \times 10^{-11}$ | 2323 | $1.70 \times 10^{-12}$ | 791.2 |
| Becavizumab: average results |  |  |  |  |  |  |
| S. L. (%) | $t_c$ (days) | $\epsilon$ | $D_{PCL_1}$ (cm<sup>2</sup>/s)* | $\tau_2$ | $D_{PCL_2}$ (cm<sup>2</sup>/s)* | Error value |
| 5 | 3 | $4.36 \times 10^{-3}$ | $1.36 \times 10^{-11}$ | 4658 | $6.70 \times 10^{-13}$ | 9.84 |
| | 7 | $3.09 \times 10^{-3}$ | $6.83 \times 10^{-12}$ | 3420 | $6.47 \times 10^{-13}$ | 19.33 |
| | 14 | $2.30 \times 10^{-3}$ | $3.79 \times 10^{-12}$ | 2616 | $6.30 \times 10^{-13}$ | 37.15 |
| 7.5 | 3 | $8.95 \times 10^{-3}$ | $5.74 \times 10^{-11}$ | 3865 | $1.66 \times 10^{-12}$ | 247.8 |
| | 7 | $6.17 \times 10^{-3}$ | $2.73 \times 10^{-11}$ | 2859 | $1.55 \times 10^{-12}$ | 389.8 |
| | 14 | $4.92 \times 10^{-3}$ | $1.73 \times 10^{-11}$ | 2667 | $1.32 \times 10^{-12}$ | 539.5 |
| 10 | 3 | $9.96 \times 10^{-3}$ | $7.10 \times 10^{-11}$ | 3281 | $2.17 \times 10^{-12}$ | 270.5 |
| | 7 | $7.03 \times 10^{-3}$ | $3.54 \times 10^{-11}$ | 2751 | $1.83 \times 10^{-12}$ | 514.3 |
| | 14 | $5.53 \times 10^{-3}$ | $2.19 \times 10^{-11}$ | 2342 | $1.69 \times 10^{-12}$ | 796.5 |

**Fig. C8.**
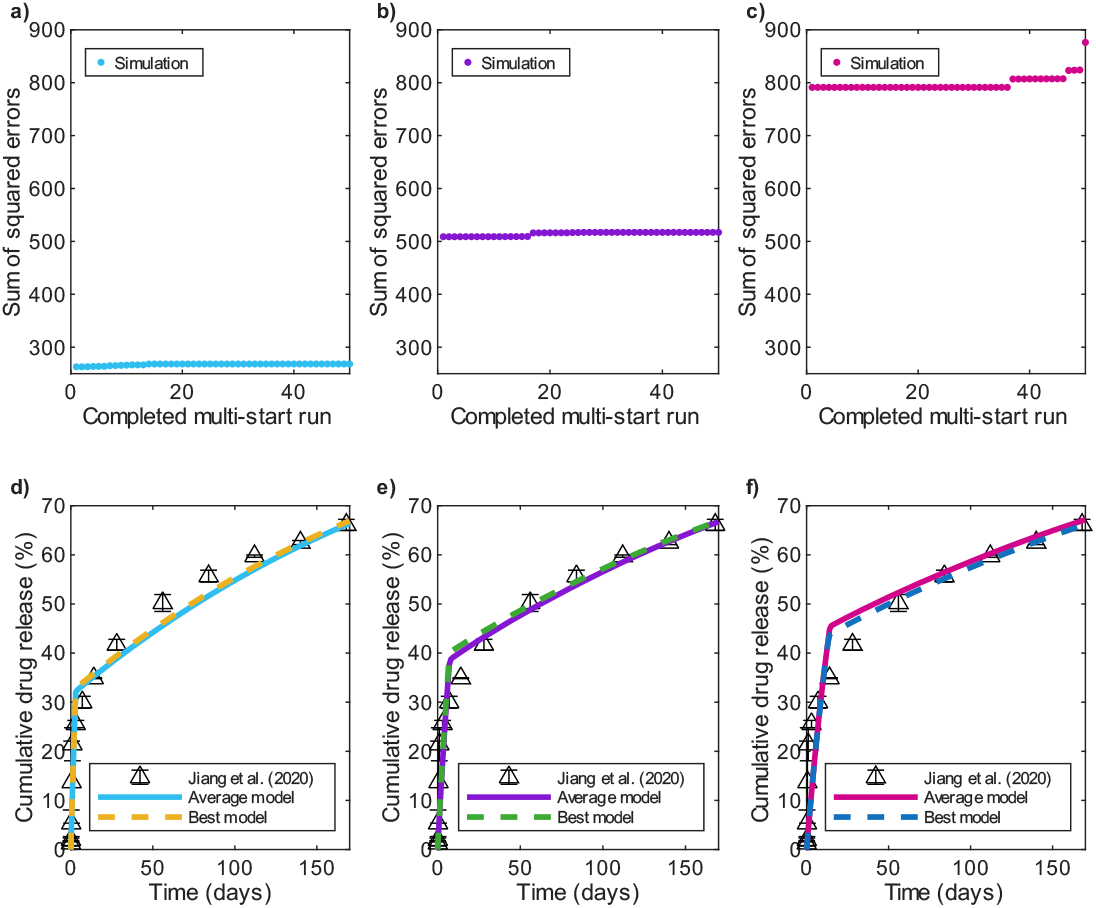
Cumulative bevacizumab release profiles after multi-start parameter estimation in 10% salt leaching PCL-only microcapsules with parameters in Table C1. First column: *t*_*c*_ = 3 days. Second column: *t*_*c*_ = 7 days. Third column: *t*_*c*_ = 14 days. First row: Error values. Second row: Average and best models compared to experimental data. Experimental data from Jiang et al. [11].

## Appendix D Model predictions at early time points

**Fig. D9.**
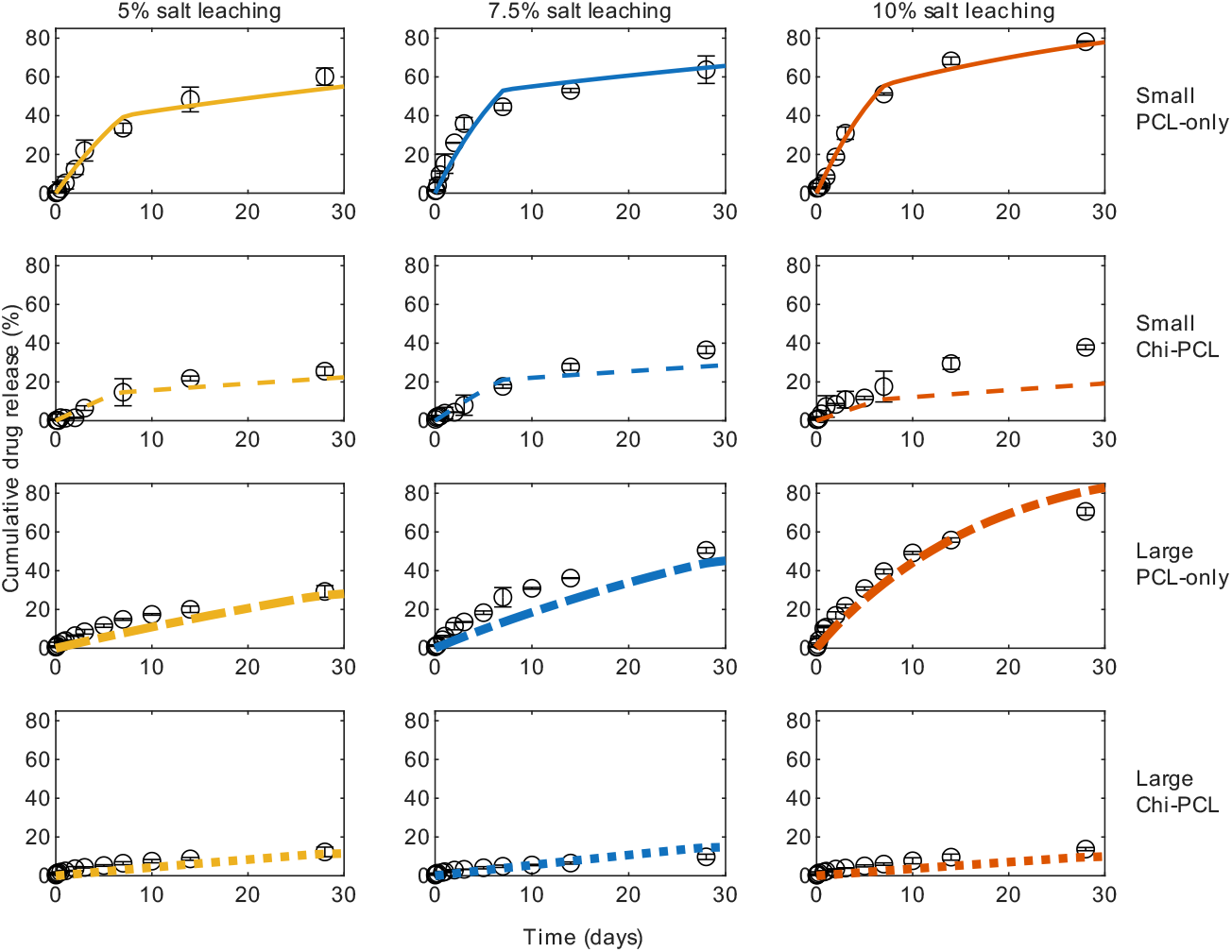
Cumulative BSA release profiles during the first 30 days. Predictions obtained after multi-start parameter estimation in different microcapsule formulations with parameters in Table 3. Experimental data from Jiang et al. [11]. BSA: bovine serum albumin. Chi: chitosan. PCL: polycaprolactone.

**Fig. D10.**
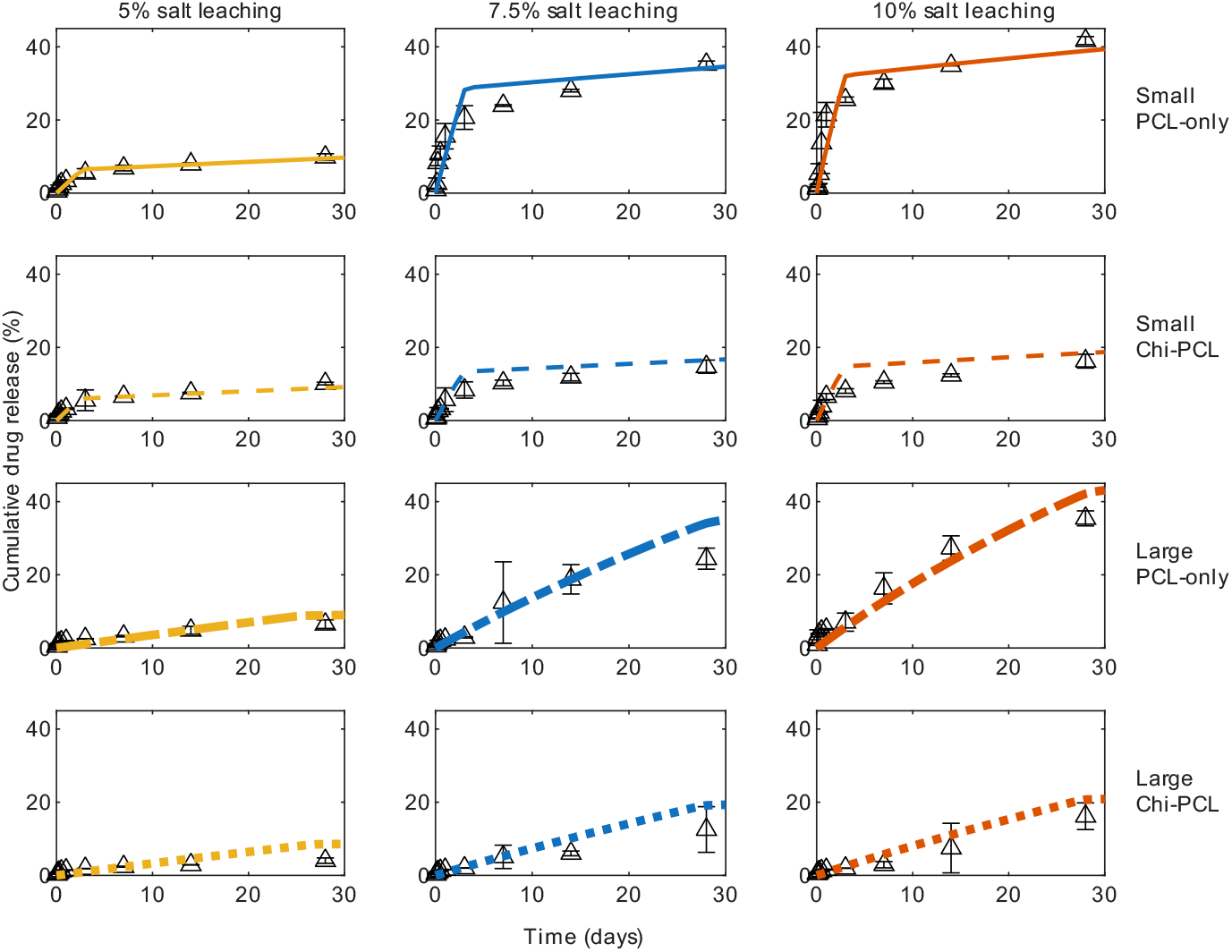
Cumulative bevacizumab release profiles during the first 30 days. Predictions obtained after multi-start parameter estimation in different microcapsule formulations with parameters in Table 4. Experimental data from Jiang et al. [11]. Chi: chitosan. PCL: polycaprolactone.

## Appendix E Cumulative release at dimensionless time from mono-layered microcapsules

The dimensionless time 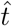 is defined as

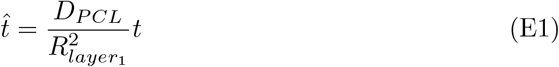

where *D*_*PCL*_ represents the drug diffusion coefficient in PCL, 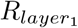 represents the PCL radius of a mono-layered microcapsule, and *t* corresponds to time. Recall Equation (19), where for the fast early release phase, 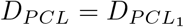 for 0 ≤ *t* ≤ *t*_*c*_. and for the slower second release phase, 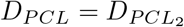 for *t > t*_*c*_.

Figure E11 shows the cumulative BSA release profiles as a function of dimensionless time for the three formulations (each column) of small and large PCL-only microcapsules. The first row in Figure E11 corresponds to the small microcapsules after parameter estimation (Table 3). The model predictions (black) reproduce the biphasic release behavior observed experimentally: the early release phase aligns closely with the experimental data (yellow), and the later phase also remains consistent with the experimental trend (blue).

When reusing the parameters derived from the small microcapsules to the large microcapsules (Figure E11, second row), the model substantially underpredicts drug release. This discrepancy suggests the presence of an additional mechanism (e.g., a partial release from the capsule caps) that was not incorporated into the initial large microcapsule model. Incorporating this mechanism and extending the critical time to *t*_*c*_ = 28 days (Figure E11, third row and Table 3) markedly improved the agreement between model predictions (purple curves) and experimental data (green and orange for late and early phases, respectively), corroborating the need of accounting for this process in the larger microcapsule.

Among the large microcapsule formulations with updated parameters, the 7.5% and 10% salt leaching formulations showed the greatest deviation from experimental data. In the 7.5% salt leaching formulation, the model fails to capture the release behavior both prior to the critical time and at later time points. For the 10% formulation, the mismatch becomes evident after the phase transition. These discrepancies may be linked to the unusually rapid release observed in the small 7.5% formulation, which likely led to higher parameter estimates. In the case of the 10% formulation, the faster release of the large microcapsules compared with the small ones appears to have influenced the parameter estimation, thereby contributing to the observed divergence.

Figure E12 presents the cumulative bevacizumab release profiles for the three formulations (each column) of small and large PCL-only microcapsules. The first row in Figure E12 corresponds to the small microcapsules following parameter estimation (Table 4). The fitted model predictions captured the bi-phasic release behavior observed experimentally, with good agreement in both the early and later phases of release.

Using the parameters obtained from the small microcapsules for the large microcapsules (Figure E12, second row), results in substantial underprediction of the experimental release. Incorporating a partial drug release from the caps to the large microcapsules model and extending the critical time to *t*_*c*_ = 28 days (Figure E12, third row and Table 4) markedly improved the agreement between model predictions and experimental data.

**Fig. E11.**
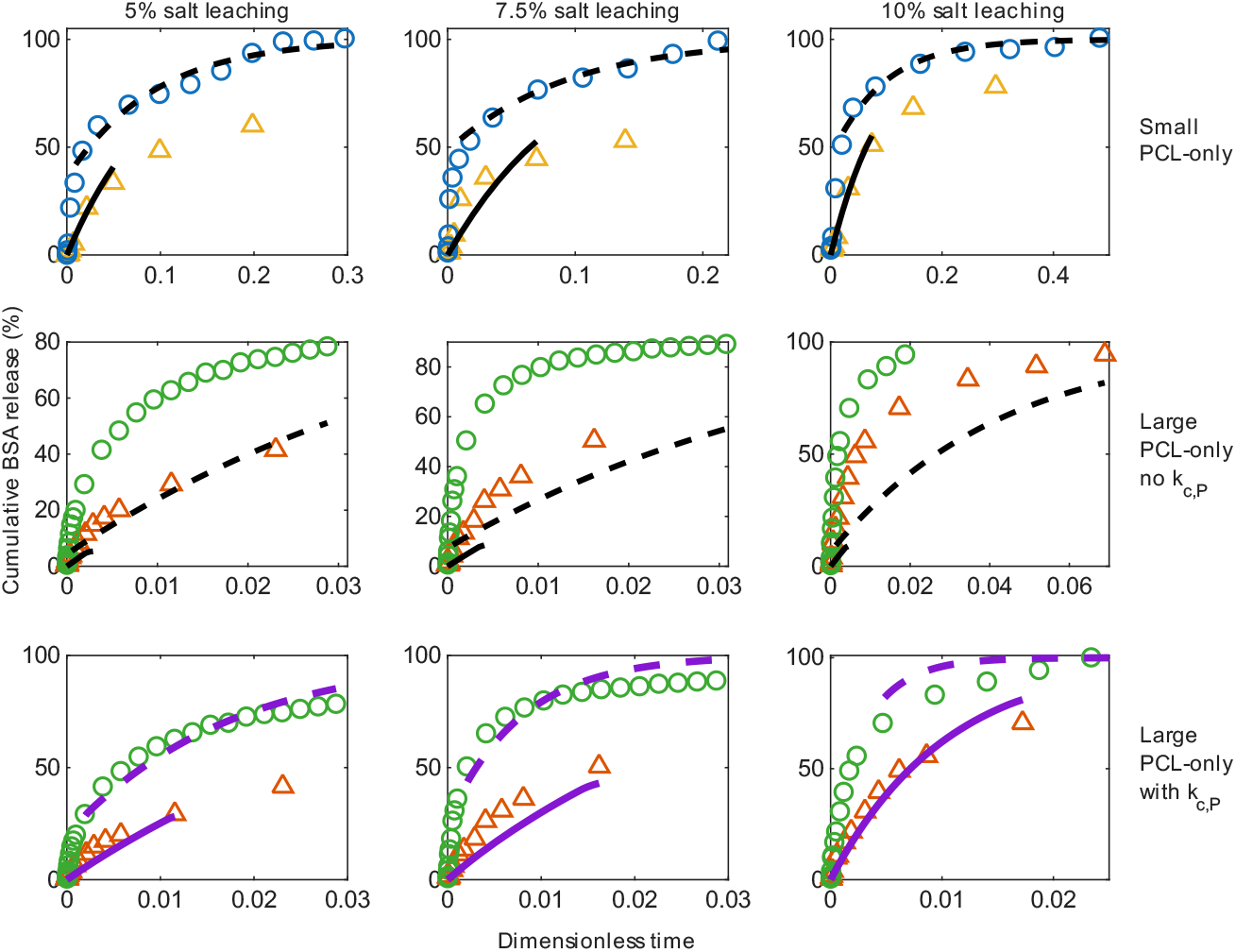
BSA cumulative release profiles after parameter estimation. First row: small microcapsules. Second row: large microcapsules using estimated parameters from small microcapsules, *t*_*c*_ = 7 days, and no flux boundary conditions at the end caps. Third row: large microcapsules using estimated porosity and tortuosity from small microcapsules, *t*_*c*_ = 28 days, and estimating mass transfer rate *k*_*c,P*_ from the end caps. Solid curves: model predictions in the first fast diffusion regime. Dashed curves: model predictions at the second slow diffusion regime. The black curves indicate using small PCL-only parameters without mass transfer from the ends, and the purple curves use parameters refitted for large PCL-only microcapsules with mass transfer from the ends. Circles: experimental data following the slow diffusion regime. Blue circles are for the small microcapsules, and green circles are for the large microcapsules. Triangles: experimental data following the fast diffusion regime. Yellow triangles are for the small microcapsules, and orange triangles are for the large microcapsules. *k*_*c,P*_ : mass transfer rate in large PCL-only microcapsules. Experimental data from Jiang et al. [11].

**Fig. E12.**
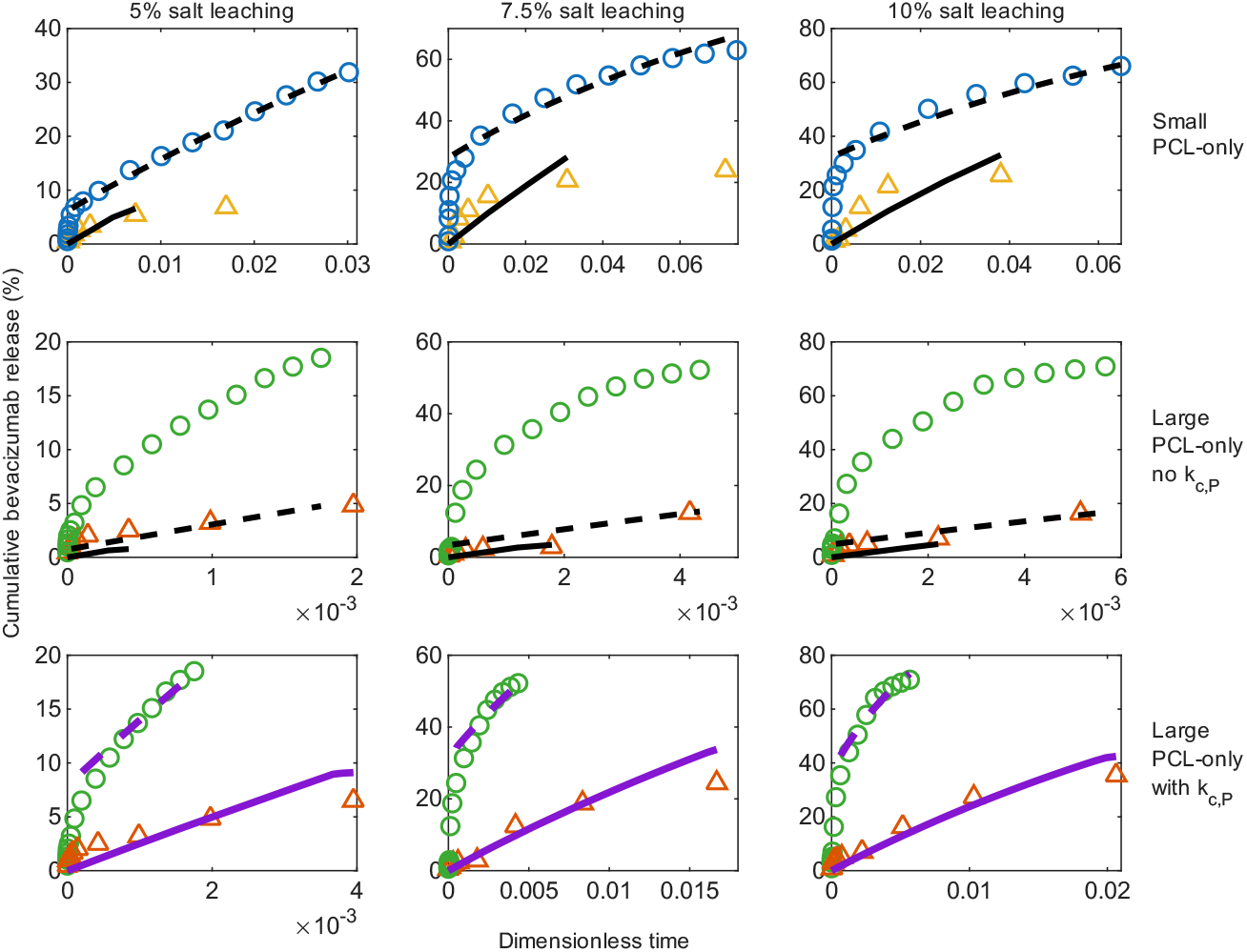
Bevacizumab cumulative release profiles after parameter estimation. First row: small microcapsules. Second row: large microcapsules using estimated parameters from small microcapsules, *t*_*c*_ = 3 days, and no flux boundary conditions at the end caps. Third row: large microcapsules using estimated porosity and tortuosity from small microcapsules, *t*_*c*_ = 28 days, and estimating mass transfer rate *k*_*c,P*_ from the end caps. Solid curves: model predictions in the first fast diffusion regime. Dashed curves: model predictions at the second slow diffusion regime. The black curves indicate using small PCL-only parameters without mass transfer from the ends, and the purple curves use parameters refitted for large PCL-only microcapsules with mass transfer from the ends. Circles: experimental data following the slow diffusion regime. Blue circles are for the small microcapsules, and green circles are for the large microcapsules. Triangles: experimental data following the fast diffusion regime. Yellow triangles are for the small microcapsules, and orange triangles are for the large microcapsules. *k*_*c,P*_ : mass transfer rate in large PCL-only microcapsules. Experimental data from Jiang et al. [11].

